# Quantification of new and archived *Diaphorina citri* transcriptome data using a chromosomal length *D. citri* genome assembly reveals the vector’s tissue-specific transcriptional response to citrus greening disease

**DOI:** 10.1101/2021.09.19.460957

**Authors:** Marina Mann, Surya Saha, Joseph M. Cicero, Marco Pitino, Kathy Moulton, Lilianna Cano, Wayne B. Hunter, Lukas A. Mueller, Michelle Heck

**Author notes:** To whom correspondence should be addressed: Michelle Heck.

## Abstract

Background

Huanglongbing (HLB) is the most serious disease of citrus. HLB is caused by the obligate, intracellular bacterium “*Candidatus* Liberibacter asiaticus” (*C*Las). *C*Las is transmitted by *Diaphorina citri*, the Asian citrus psyllid. Development of transmission blocking strategies to manage HLB relies on knowledge of *C*Las-*D. citri* interactions at the molecular level. Prior transcriptome analyses of *C*Las-infected and un-infected *D. citri* point to changes in psyllid biology due to *C*Las-infection. These studies relied on incomplete versions of the *D. citri* genome, lacked proper host plant controls, and/or were analyzed using different statistical approaches. Therefore, we used standardized experimental and computational approaches to identify differentially expressed genes in both CLas (+) and CLas (-) *D.* citri. The comparative analysis utilized the newest chromosomal length *D. citri* genome assembly Diaci_v3. In this work, we present a quantitative transcriptome analysis of excised heads, salivary glands, midguts and bacteriomes from *C*Las (+) and *C*Las (-) insects.

Results

Each organ had unique transcriptome profiles and responses to *C*Las infection. Though most psyllids were infected with *C*Las, *C*Las-derived transcripts were not detected in all organs. By analyzing the midgut dataset using both the Diaci_v1.1 and v3.0 *D. citri* genomes, we showed that improved genome assembly led to significant and quantifiable differences in RNAseq data interpretation.

Conclusions

Our results support the hypothesis that future transcriptome studies on circulative, vector-borne pathogens should be conducted at the tissue specific level using complete, chromosomal-length genome assemblies for the most accurate understanding of pathogen-induced changes in vector gene expression.

## Background

Huanglongbing (HLB), also known as citrus greening, is the most serious disease of citrus (reviewed in [1–3]). HLB symptoms include leaves with blotchy chlorotic mottling, stunting, loss of root biomass, premature fruit drop, uneven fruit development, and ultimately tree death. HLB is associated with plant vascular tissue infection by the gram-negative, uncultivable alpha-proteobacteria “*Candidatus* Liberibacter asiaticus” (*C*Las), “*Ca*. L. americanus” (*C*Lam) and “*Ca*. L. africanus” (*C*Laf). The Asian citrus psyllid *Diaphorina citri* Kuwayama (Hemiptera: Liviidae) is the vector of *C*Las and *C*Lam, whereas the African citrus psyllid *Trioza erytreae* (Del Guercio) is the vector of *C*Laf. HLB is found in most regions where citrus is cultivated, including in the United States where it has decimated a multi-billion dollar industry in Florida and is threatening the industries in Texas and California [4]. HLB affects all genotypes of *Citrus* and some other members of Rutaceae. Liberibacter, like other vascular plant pathogens, are also readily transmitted from plant to plant by grafting [5, 6], a technique which puts phloem from the vascular tissue of one plant into contact with that of another. However, psyllid transmission remains the primary driver of HLB epidemiology in citrus groves.

Evidence thus far on *C*Las transmission by *D. citri* is consistent with a circulative, propagative transmission mode that is inextricably linked to the insect’s development and intracellular environment surrounding *C*Las bacteria (**Figure 1**) [7]. During the circulative propagative transmission cycle of *C*Las, *D. citri* acquire *C*Las from an infected citrus host during phloem ingestion as early as the 2^nd^ nymphal instar [8] but in increasing amounts during the 4^th^ and 5^th^ instars of the nymphal stage [9]. The bacteria remain associated with the insect during molting [9, 10]. *C*Las circulates throughout the body of *D. citri* until it reaches the salivary gland tissues, where it replicates to high levels in the adults [11–14]. The titer of *C*Las increases, presumably in the salivary gland tissue, over approximately 1-2 weeks [9]. The infected adults inoculate the bacteria back into the same tree, or in the case of facilitating spread, a different tree and complete the transmission cycle. *C*Las can be found in cells of the insect’s alimentary canal, especially the midgut [11, 12, 15]. The bacteria also systemically infect the psyllid during propagative transmission, including the hemolymph, salivary glands, muscle, fat body and reproductive organs (reviewed in [3]). Specific cellular receptors in these different *D. citri* tissues are not known. In adults, *C*Las forms a biofilm along the midgut and induces apoptosis of midgut epithelial cells [16], a process which is not observed in nymph midguts [17]. In the midgut, the bacterium is hypothesized to be associated with the endoplasmic reticulum based on microscopic observations [18].

**Figure 1.**
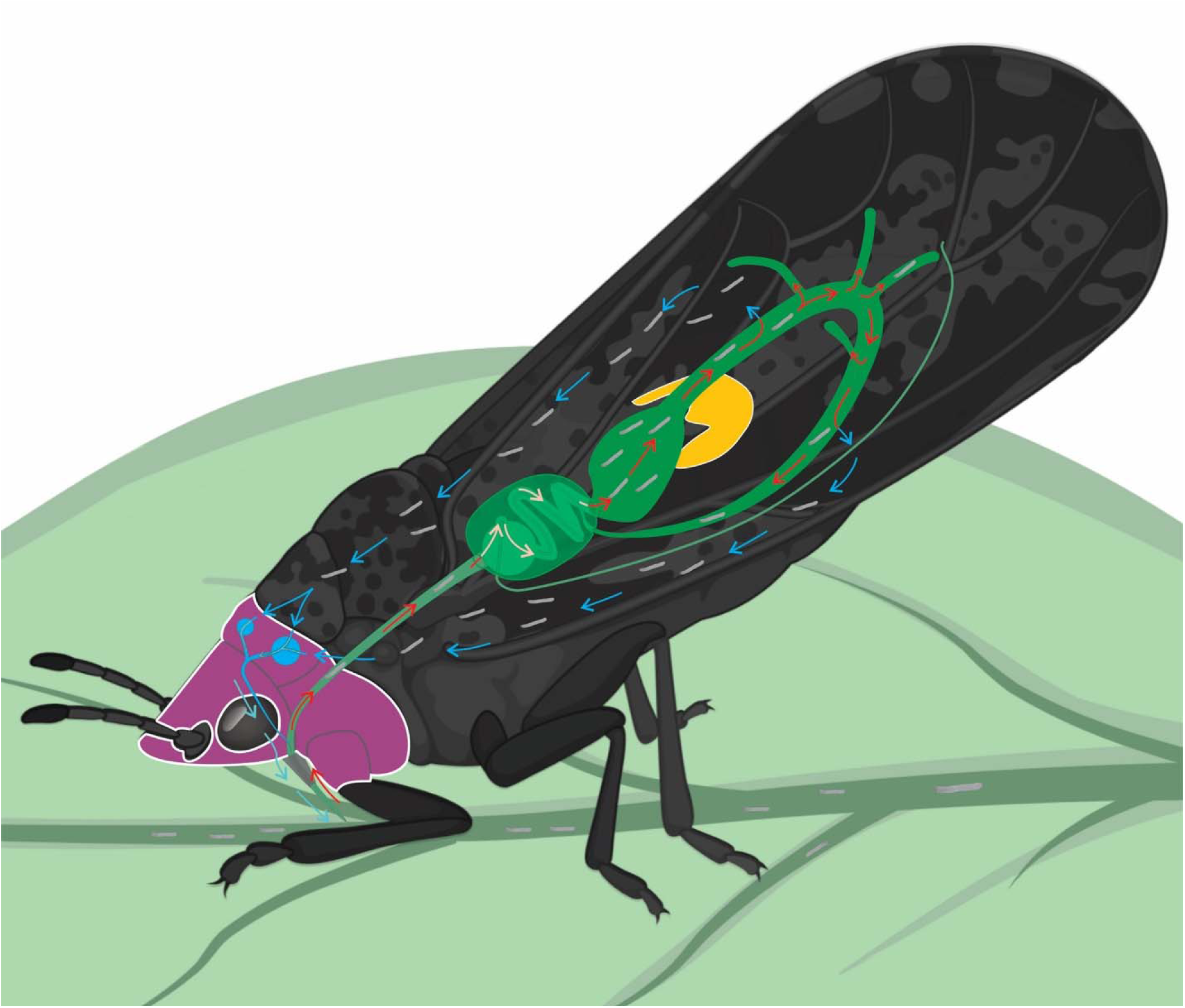
Schematic of *Diaphorina citri* on a citrus leaf, showing the anatomical location and physical details of four parts that were extracted from adult *D. citri* to create four datasets (gut - green, bacteriome - yellow, salivary gland - blue, head – dark purple). The circulative transmission of “*Candidatus* Liberibacter asiaticus” (*C*Las represented by small grey lines) is represented as *C*Las travels from leaf veins through the gut, crossing the midgut epithelial cell layer to circulate in the body of *D. citri* where it is known to enter and possibly interact with many different organs. *C*Las enters the salivary gland where it is known by contributory effects from acquisition by late instar nymphs, to replicate to high levels, at which point it can be inoculated into the phloem while adult *D. citri* feed (see 3D imaging and digital video by Alba-Tercedor et al. (2021) for more details [42]). *C*Las (-) adults transmit *C*Las inefficiently if the bacteria are acquired during the adult stage.

Not all psyllids become infected with *C*Las when feeding on *C*Las-infected trees, and not all psyllids transmit efficiently even if infected [10, 19, 20]. Such variability has undoubtedly hampered the ability to disentangle transcriptomic and proteomic responses in psyllids from the effects of *C*Las directly or indirectly, and from the effect of *C*Las-infected trees. Additionally, different psyllid populations vary in their ability to acquire and transmit the bacterium [10]. These variable acquisition and transmission rate efficiencies are heritable traits in *D. citri*, supporting the idea that *D. citri* genes and proteins at least partially regulate the ability to transmit *C*Las. Due to the variability in *D. citri* infection with *C*Las, in ‘omics and systems biology studies on *D. citri* when measurements are based on responses of hundreds or thousands of pooled insects or insect tissues, insects reared on *C*Las-infected trees are referred to as “*C*Las- exposed insects” or “*C*Las-infected” and *D. citri* reared on healthy citrus as “non-exposed” or “non-infected”. Other papers referred to these groups of insects more generally as *C*Las (+) and *C*Las (-) insects to denote the infection state of the insect population as a whole, since not all individuals reared on an infected tree will become infected with *C*Las. This paper will use the *C*Las (+) and *C*Las (-) designation to refer to the different sample groups, where the *C*Las (+) insects were reared on *C*Las-infected trees and the *C*Las (-) insects were reared on healthy citrus which also tested negative for *C*Las by quantitative PCR (qPCR).

*D. citri* harbors three bacterial symbionts, “*Candidatus* Profftella armatura,” “*Candidatus* Carsonella ruddii,” and *Wolbachia-Diaphorina* (wDi) [21–27] which reside in a specialized organ referred to as the bacteriome. The bacteriome is comprised of bacteriocytes – psyllid cells densely packed with the endosymbiotic bacteria. The bacteriome of *D. citri* has a precise and elegant cellular organization that has been described using fluorescence microscopy [21, 25]. *Carsonella* resides in the outer bacteriocytes and *Profftella* resides in the internal syncytial cytoplasm of the bacteriome. The function of these beneficial bacterial symbionts in the biology of *D. citri* is inferred from bacterial genome sequencing, proteomics and metabolite data. Evidence shows that these symbionts have complex and possibly shared, coordinated metabolic and protein signaling networks with *C*Las [28, 29]. While no direct evidence exists to support a role for the bacterial symbionts in *C*Las acquisition and transmission, *Profftella* has been shown to modulate production of diaphorin, a polyketide encoded in the *Profftella* genome, in response to *C*Las [29, 30].

Rapid advances in genome sequencing technologies have paved the way to a deeper understanding of vector biology over the past decade, including in the analysis of the *D. citri* genome sequence [23, 31–34]. The short read-based assembly, Diaci_v1.1 [33, 35], has been foundational to the vast majority of published research on *D. citri* to date, including the newest chromosomal length reference genome [34], which is expected to lend more reliability and contain more cohesive, full-length annotated gene models. Numerous studies have used these valuable *D. citri* genome sequencing resources to investigate interactions between *D. citri* and *C*Las and *D. citri* biology at the transcriptome and proteome levels [15, 29, 36–38]. Wu and colleagues [39] published a thorough RNAseq experiment including an analysis of organs, sexes, and life stages of *D. citri*. Their analysis focused on potential insecticide detoxification genes from *C*Las (-) insects raised on a close relative of citrus known to be resistant to systemic infection by *C*Las, *Murraya exotica*, but did not address the impact of *C*Las infection in these organs. A year later, the same group published a paired transcriptome-proteome paper focusing on *C*Las (-) *D. citri* salivary glands and associated salivary secretions [40]. They focused on identifying bioactive molecules from the saliva and salivary gland ‘omics analysis and discussed proteins that were found uniquely in the salivary glands from *D. citri* reared on healthy plants.

Tissue-specific omics analyses enables a molecular snapshot of *C*Las-*D. citri* interactions within specific tissues known to be colonized by *C*Las in the insect. Studies have revealed stark differences in patterns of expression when comparing tissue-specific responses to whole body responses [15, 38]. However, earlier studies were limited in the interpretation of the data because of the incomplete nature of the *D. citri* genome that were used as a backbone for the quantitative analysis and the application of different computational workflows to identify differentially expressed genes. Kruse et al. (2016) did a thorough analysis and discussed the midgut transcriptomics responses to *C*Las using four biological replicates of pools of hundreds of midguts and performed dual differential expression analysis using two types of computational biology tools, edgeR and DESeq2, to reduce the false discovery rates [15]. However, the results were dependent on paired proteomics and transcriptomics that were both aligned to the relatively low quality and incomplete v1.1 *D. citri* genome, the assembly available at the time. Yu and colleagues [41] built on the Kruse et al. study [15] using the *D. citri* v2.0 genome, which also lacked the Hi-C scaffolding included in the newest v3.0 genome. Despite the limitations of the genome sequences used for the analyses of these transcriptomes, the results clearly showed that *C*Las has different effects on metabolic pathways expressed within different tissues of *D. citri*. To understand the nature of the *C*Las-*D. citri* relationship at the molecular level, a holistic approach which both integrates the responses across different tissues involved in the circulative transmission pathway and quantifies the impact of *C*Las infection on the transcriptional regulation within specific tissues is necessary.

In this work, we compare *C*Las (-) to *C*Las (+) psyllid datasets from four different organs: excised *D. citri* midguts, bacteriomes, salivary glands, and heads, using the newest *D. citri* genome assembly (v3.0), which includes chromosomal length scaffolds [34]. This study advances our understanding of *D. citri*-*C*Las interactions because it integrates an analysis of new transcriptome data with previously published transcriptome data to show the impact of *C*Las on the transcriptional landscape of *D. citri* organs involved in the circulative, propagative transmission. The difficulties of comparing the four datasets – three of which were collected from separate insect colonies, at different times, sequenced separately, stored in freezers for different lengths of time, and contain variable amounts of *C*Las in each tissue type – should be acknowledged. This study does not purport to have controlled for all differences found between these datasets, but we do attempt to carefully explain results within the bounds of our controls and include caveats for the confounding effects. Our analysis demonstrates that it is possible to analyze new ‘omics data in the context of and alongside historical data in public repositories to maximize the use of existing large-scale dataset resources in discovering new biology. The results underscore the importance of chromosomal length assemblies of arthropod genomes for accurate interpretation of gene expression.

## Data Description

### Experimental design, RNA collection, and sequencing of four *D. citri* RNA datasets

Psyllid colonies and citrus plants used to generate samples for the bacteriome, head and midgut datasets were continuously maintained by the USDA ARS in Ithaca NY and the USDA ARS in Fort Pierce, FL under the same growth conditions. These psyllid colonies – including *C*Las (-) and *C*Las (+) *Diaphorina citri* adults and nymphs raised on *Citrus medica* (Citron) – were originally started in 1999 from individuals collected from a farm near Fort Pierce, Florida and the *C*Las strain used came with those original individuals. Growth chambers were maintained at 22.8°C-26.7°C, 70-80% humidity and a 14h light/10h dark photoperiod. Citrus plants (*Citrus medica*, Citron) were grown in greenhouse conditions from seed. *C*Las (+) *C. medica* were inoculated using *C*Las (+) *D. citri*. When insect colonies contained 1-2 week old adults, pools of adult *D. citri* were collected from each colony to create each biological replicate (120 per bacteriome and head replicate (Ithaca colony), 150 per salivary gland replicate (Fort Pierce colony), 250 per midgut replicate (Fort Pierce colony described in [15]). Insects were anesthetized on ice for a few hours prior to and during dissection.

#### Bacteriome and head samples

Using a dissecting scope, bacteriomes and heads of adult psyllids were excised into mili- Q (MQ)-water then moved to 2ml tubes containing 350ul of buffer RLT (Qiagen RNeasy kit) with beta-mercaptoethanol and kept on ice during collections. Once the collection of a biological replicate was complete, the tubes containing pools of psyllid organs were flash frozen in liquid nitrogen and stored in -80°C until needed. Total RNA was extracted following the Qiagen RNeasy extraction protocol, including sample disruption with syringes and DNase treatment to remove DNA contamination.

#### Salivary glands and midguts

Salivary tissues and midguts were preserved in TriZol. Salivary glands were excised as described by Cicero and Brown [43] in pools of 300 per replicate in TRIzol LS (ThermoFisher). Samples were kept at -80 ° C (bioreps 1-3, *C*Las (-/+) were kept 1 year, while replicate 4, both *C*Las (-/+), was kept for 2 years) prior to RNA extraction. Total RNA was extracted for both midguts and salivary glands following the standard TRIzol RNA extraction protocol [44] including light syringe disruption prior to adding ethanol, and DNase treatment to purify total RNA. Total RNA quality was tested using an RNA gel prior to library preparation. Details of midgut sample handling can be found in Kruse et al. [15].

Illumina libraries for all samples were made by Polar Genomics LLC following the protocol of Zhong et al., [45] and included poly-A tailed mRNA enrichment. Libraries were shipped on dry ice to GENEWIZ where they were pooled for Illumina paired-end 150bp sequencing. Bacteriome, head and salivary gland samples were sequenced separately from the previously published midgut samples. Raw data has been uploaded to NCBI and is accessible to reviewers via BioProject accession # PRJNA385527, submission ID SUB10382129 and will be made available to the public upon publication.

## Analyses

### Though most psyllids were infected with *C*Las, *C*Las-derived RNAseq reads were not detected in all organs

Using quantitative PCR (pPCR) analysis of whole insects, we determined the *C*Las infection rate of the *D. citri* populations used to generate samples of each organ, which informed our interpretations of differential expression and infection for the subsequent datasets. The *C*Las infection rate was for all samples were derived from sampling whole insects, not dissected organs. Across all sample types, the percent infection rate ranged between 73-85%. Cq values lower than 40 were counted as *C*Las (+) (Table 1). In addition to a population-level assessment of *C*Las infection, we quantified *C*Las-mapped reads found within each sample after sequencing (Table 1 and Figure S1). Read counts mapping to the *C*Las-psy62 genome (genome produced from a single psyllid in FL [46]) were significantly detected in *C*Las (+) salivary gland and head samples (an average of 1965 and 2681 reads, respectively). This might be influenced by the fact that only AT-rich transcripts from the CLas-psy62 genome were captured during the poly-A enrichment step, prior to library prep. Upon closer analysis of the *C*Las-aligning reads from the salivary glands, when at least three biological replicates had a transcript with at least one read, 50 different *C*Las transcripts were represented, with an additional six rRNA transcripts (three of each 16S and 23S transcripts), for a total of 56 *C*Las-psy62 transcripts identified. The majority of *C*Las reads from the salivary glands aligned to the top 10 transcripts, where the total number of reads across all biological replicates of each transcript ranged from 80 to 290. Of these top 10, three were listed as “protein coding” and annotated as figB, figC, and parB, while the rest were unlabeled/unknown (Table S1). While these numbers are not enough to allow for statistical analysis, they present an intriguing picture of *C*Las infection in the organ essential for successful transmission.

**Table 1.**
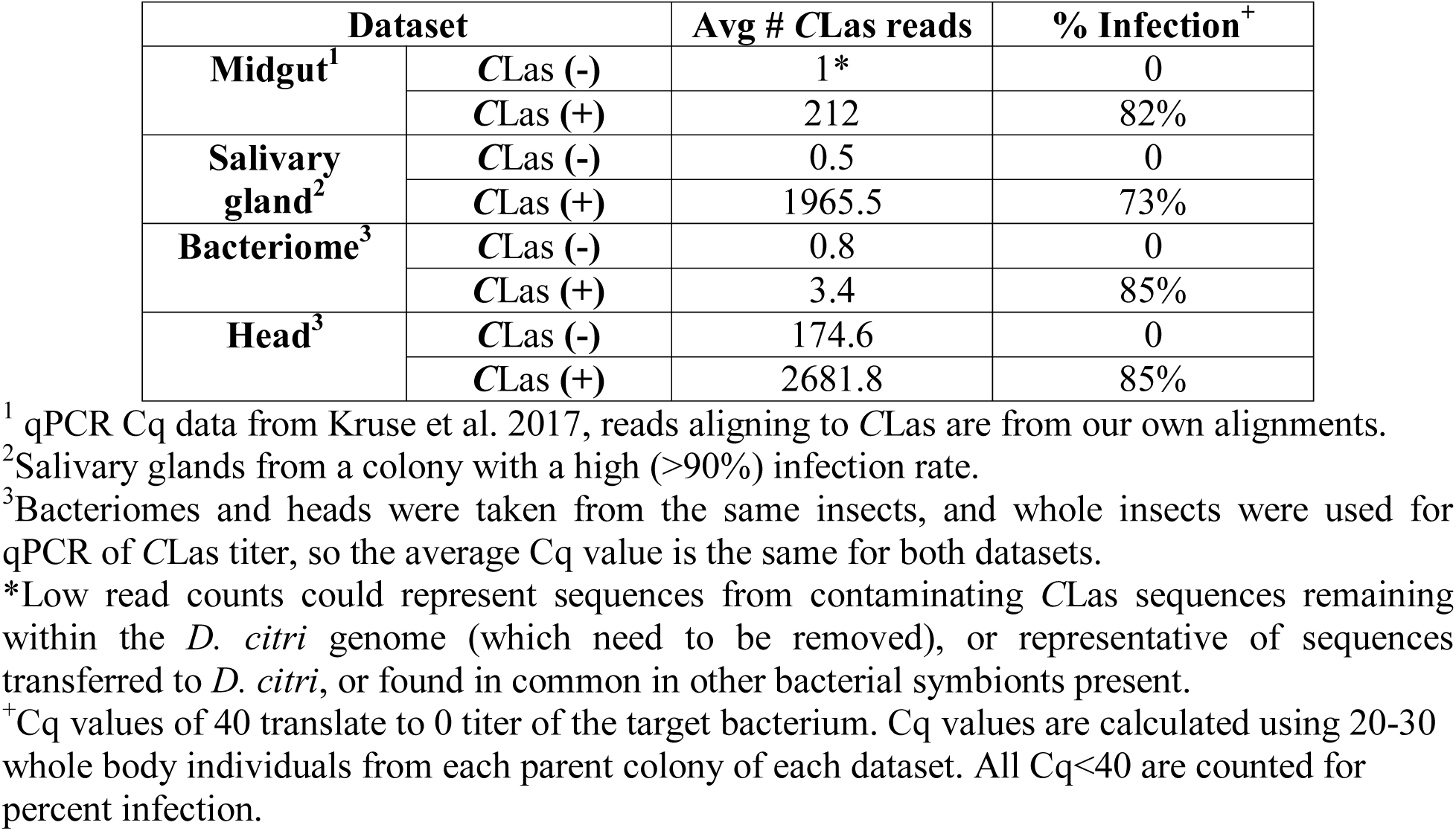
Percent infection by *C*Las in different *Diaphorina citri* tissues as measured by qPCR, and the average number of RNAseq reads that aligned to the *C*Las genome (psy62) from each dataset.

| Dataset |  | Avg # CLas reads | % Infection<sup>+</sup> |
| --- | --- | --- | --- |
| Midgut<sup>1</sup> | CLas (-) | 1* | 0 |
|  | CLas (+) | 212 | 82% |
| Salivary gland<sup>2</sup> | CLas (-) | 0.5 | 0 |
|  | CLas (+) | 1965.5 | 73% |
| Bacteriome<sup>3</sup> | CLas (-) | 0.8 | 0 |
|  | CLas (+) | 3.4 | 85% |
| Head<sup>3</sup> | CLas (-) | 174.6 | 0 |
|  | CLas (+) | 2681.8 | 85% |

<sup>1</sup> qPCR Cq data from Kruse et al. 2017, reads aligning to CLas are from our own alignments.
<sup>2</sup>Salivary glands from a colony with a high (>90%) infection rate.
<sup>3</sup>Bacteriomes and heads were taken from the same insects, and whole insects were used for qPCR of CLas titer, so the average Cq value is the same for both datasets.
\*Low read counts could represent sequences from contaminating CLas sequences remaining within the *D. citri* genome (which need to be removed), or representative of sequences transferred to *D. citri*, or found in common in other bacterial symbionts present.
<sup>+</sup>Cq values of 40 translate to 0 titer of the target bacterium. Cq values are calculated using 20-30 whole body individuals from each parent colony of each dataset. All Cq<40 are counted for percent infection.

### Global assessment of four transcriptomics datasets clarifies the *D. citri* organ-specific response to *C*Las

Across all four datasets, we obtained an average of 27.23 million high-quality reads, (midguts: 26.43M, salivary glands: 44.98M, bacteriomes: 22.11M, heads: 15.40M), and 71.3% of the reads aligned concordantly to the v3.0 *D. citri* genome on average (average concordant alignment in midguts: 74.17%, salivary glands: 73.51%, bacteriomes: 81.12%, heads: 56.43). The head dataset proved to be more variable as compared to the other datasets, recording the least number of raw reads and the lowest average percent alignment. In contrast, the highest percent alignment to the *D. citri* genome was recorded by the bacteriome dataset, samples of which were collected from the same individual insects as the head dataset (Table S2).

A principal components analysis (PCA) to examine the sources of variation among the four *D. citri* dataset expression profiles was performed, where each dataset includes both *C*Las (+) and *C*Las (-) biological replicates. Each organ separated from the other organs in PCA space, showing that each organ has a unique transcriptome profile. The largest source of variation (PC1 = 21%) was explained by differences in the transcriptome profiles of the midgut and bacteriome as compared to the salivary gland and head (Figure 2). The second largest source of variation between the four datasets (PC2 = 18%) was explained by differences between the midgut and the bacteriome datasets, with a smaller amount of variation between those samples and the head and salivary gland datasets along the same principal component. Importantly, biological replicates of each dataset clustered together and separately from the others (Figure S2), supporting the hypothesis that each organ has a unique transcriptomic signature independent of *C*Las infection. A closer examination of the four clusters showed that the salivary gland, bacteriome and head datasets did not differentiate between *C*Las (-) and *C*Las (+) biological replicates (Figure S2B, S2C, S2D), while midguts (Figure S2A) showed a clear separation along PC1 between *C*Las (-) and *C*Las (+) biological replicates.

**Figure 2:**
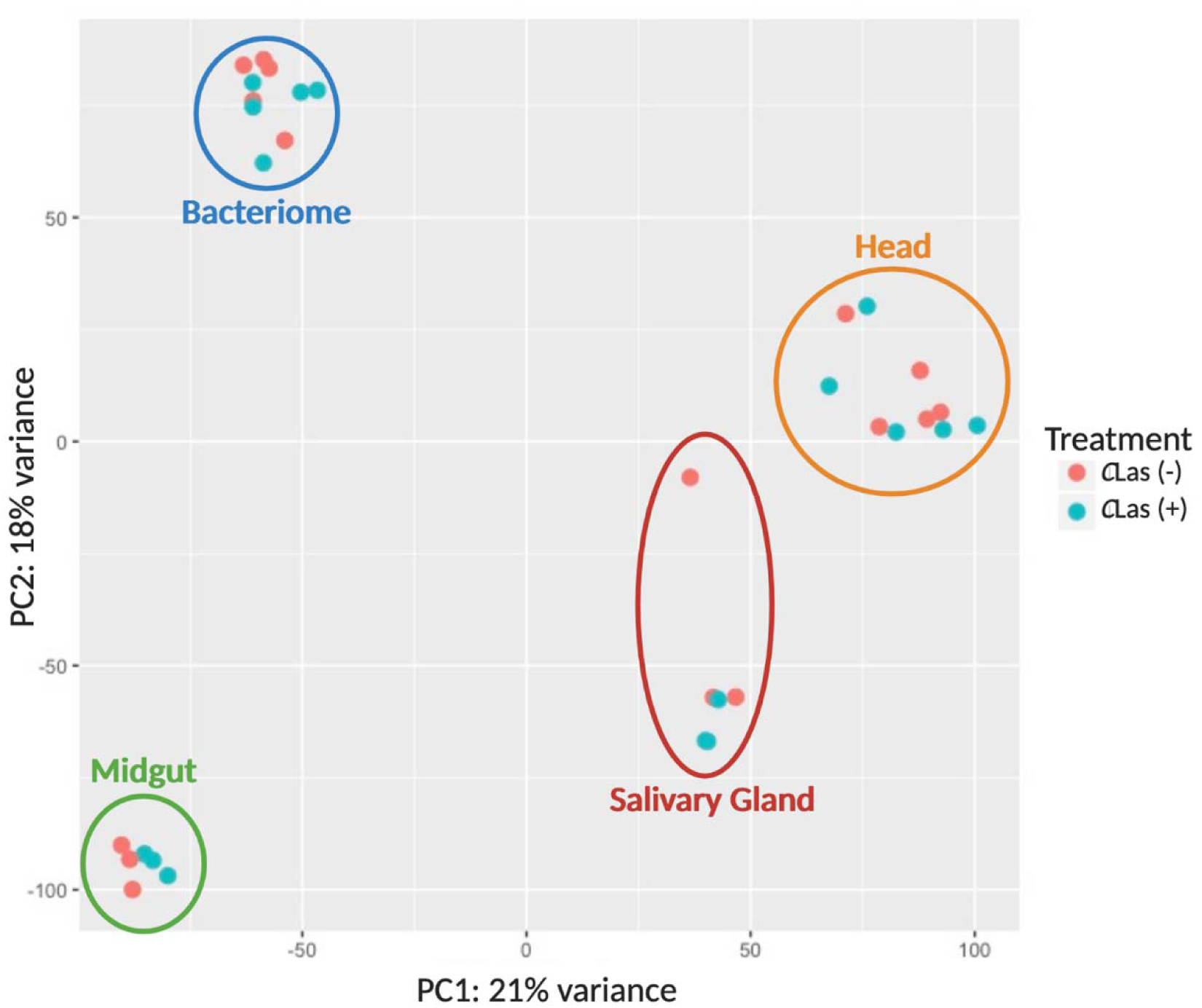
Principal components analysis (PCA) of four *Diaphorina citri* mRNAseq datasets (head, midgut, salivary gland and bacteriome), each composed of *C*Las (+) and *C*Las (-) biological replicates, showing the two main sources of variation among them. PC1 (21%) separates samples containing salivary tissues (head and salivary gland samples) from the other datasets, while PC2 (18%) distinguishes the bacteriome and head datasets (which were collected in parallel from the same individual insects), from the salivary gland and midgut datasets (which were collected independently). Within each dataset, there is little to no clear separation between *C*Las (+) and *C*Las (-) biological replicates. The remaining variance in the data, (61%) is explained by other factors in the data. Raw read counts were processed by DESeq2 using the Benjamini-Hochberg normalization method before generating the principal components plot.

PCA plots of each organ dataset comparing *C*Las (+) to *C*Las (-), revealed other sources of variation (Figure S2). The variance described by PC1 of the salivary gland dataset (44.1%, Figure S2B) was due to two samples which were kept in the -80C freezer and then sequenced a year after the other six samples, while PC2 (19%, Figure S2B) represented the effect of *C*Las infection which is not distinct, except for the two outlier samples. The bacteriome dataset (Figure S2C) showed some separation between *C*Las (+) and *C*Las (-) biological replicates (PC2=15.9%) but the majority of variation was due to variance among individual biological replicates (PC1=16.7%). The head dataset (Figure S2D) showed similar variation across all samples as the bacteriome dataset. This variation explained both the first and second major sources of variance (PC1=39.7%, PC2=27.3%) with no obvious distinctions between *C*Las (+) and *C*Las (-) biological replicates.

### Gene expression signatures in response to *C*Las infection are tissue-specific in *D. citri*

The total number of differentially expressed transcripts (including both the transcripts that were only expressed in *C*Las (+) or *C*Las (-) replicates, and the transcripts that were present but differentially expressed between *C*Las(+) and *C*Las(-) biological replicates) was determined using the maximum adjusted p-value of 0.05, yielding significantly differentially expressed transcripts in each dataset (midgut=277, salivary gland=107, bacteriome=296, head=10). From these top transcripts, those with a Log2FoldChange (L2FC) of >|2| were used for downstream analyses. This strict quality and DE threshold limited the number of final transcripts to a small number (midgut=196, salivary gland=105, bacteriome=113, head=10) (see Tables S3, S4, S5, and S6 for the list of transcripts). A skew towards up-regulated transcripts in *C*Las (+) biological replicates was detected in all organs (salivary gland: up-regulated=91, down-regulated=14; midgut: up-regulated=129, down-regulated=67; bacteriome: up-regulated=70, down-regulated=43; head: up-regulated=6, down-regulated=4).

Four major groups of transcripts were chosen based on their strong representation among the top differentially expressed gene (DEG) lists from the salivary gland, bacteriome and midgut datasets (Figure 3, Table S7) for a more detailed analysis to highlight the tissue-specific patterns of transcriptional activation in response to *C*Las. The four groups include ribosomal transcripts, immunity-related transcripts, endocytosis-related transcripts and ubiquination-related transcripts. Each dataset varies in its strength of response (as measured by L2FC and the relative number of transcripts found in each of the four categories).

**Figure 3:**
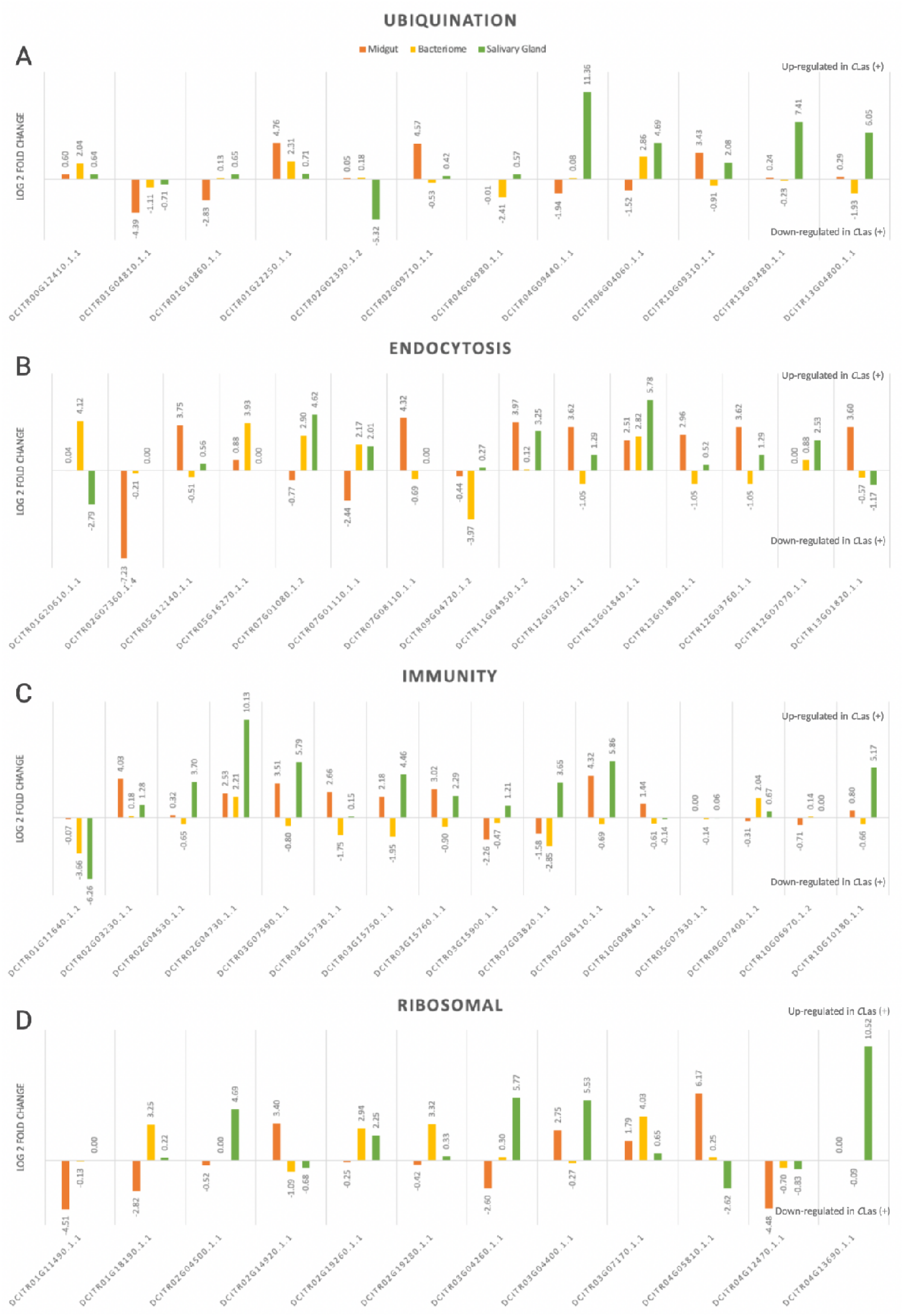
Transcripts have unique expressions across different organs of *D. citri.* The top differentially expressed (DE) transcripts from each dataset (bacteriome, midgut and salivary gland) are sorted by major functional groups including ubiquination, endocytosis, immunity and ribosomal-related transcripts. Not all transcripts are statistically DE, one transcript may be DE in one dataset, but not the others. See Table S7 for specific p-values.

Ribosomal transcript depletion in silico is known to reduce the bias of overabundant host transcripts over secondary target organism transcripts in a dataset [47, 48]. Analysis of ribosomal RNAs remaining after bioinformatic filtering left only the most highly differentially expressed rRNA transcripts between *C*Las (+) and *C*Las (-) samples. The majority of ribosomal transcripts are up-regulated in *C*Las (+) samples. Ubiquitination genes are highly upregulated in the salivary gland dataset (Figure 3A). Endocytosis genes are highly upregulated in all tissue datasets. In contrast, immunity genes are upregulated in the salivary glands and midguts but not the bacteriomes (Figure 3C). Different ribosomal genes are upregulated in all three datasets (Figure 3C).

In addition to the major patterns (Figure 3), selected transcripts of interest also showed notable changes in expression in the datasets consistent with the functions of these tissues in *D. citri* physiology that may give insight into how *C*Las is interacting with these specific tissues at the molecular level. These changes are discussed here.

#### Midgut

In addition to the organization shown in Figure 3, the top differentially expressed transcripts from the midgut dataset were manually sorted into five additional functional categories including biosynthesis and catabolism (n=55, 40 up-regulated in *C*Las (+), 15 down-regulated), cell structure and signaling (n=66, 38 up, 28 down), stress (n=10, 6 up, 4 down), transport (n=28, 19 up, 9 down), and unknown (n=37, 26 up, 11 down). The full list can be found in Table S3. Differentially expressed transcripts in the stress category include heat shock and cold shock protein genes, thioredoxin, and E3 ubiquitin ligase. Three heat shock proteins (70-A1, 70-B, 70) are up-regulated with exposure to *C*Las, while the cold shock protein is down regulated. An E3 ubiquitin ligase, a type IV collagenase and tumor protein p53 are also up-regulated. A thioredoxin transcript and a HSP20-like chaperone transcript are down-regulated with exposure to *C*Las. Transport-related transcripts that are up-regulated with *C*Las-infection include two odorant-binding protein transcripts, membrane-associated ion transporters (aquaporin, major facilitator, protein-coupled AA-transporter, efflux system protein transcript, phosphate transporter, potassium channel protein transcript, and general secretion pathway transcripts), a vacuolar-sorting protein transcript, and an intraflagellar transport particle protein transcript, among a few others. Down-regulated transcripts include syntaxin, ubiquinol cytochrome-c, membrane-associated proteins and transporters, and nuclear transport factor 2.

#### Salivary gland

The full list of statistically significant (padj<0.05) salivary gland differentially expressed (L2FC>|2|) transcripts can be found in Table S4. Transcripts for 40S and 60S subunits of the eukaryotic ribosome are highly up-regulated (40S S15a L2FC=10.12, 40S S28 L2FC=10.52, 60S L2FC=5.53), as well as six transcripts involved with transportation which are all up-regulated (ABC transporter C family L2FC=5.95, alpha-tocopherol transfer protein L2FC=8.04, gamma-glutamylcyclotransferase L2FC=8.15, geranylgeranyl transferase L2FC=6.22, MFS-type transporter L2FC=4.03, and phosphate acetyltransferase L2FC=9.30). Additionally, four elongation factor (EF) transcripts are highly up-regulated (EF-1b, EF-2, EF-4 and a Calcium-binding EF hand), consistent with increased ribosomal activity. While ubiquination-related transcripts are present in every dataset, in the salivary gland dataset two transcripts are highly up-regulated including a ubiquitin conjugating enzyme (L2FC=3.70) and ubiquitin-ligase E3 (L2FC=4.69). [41].

Since the salivary gland is known as a secretory organ, the most abundant transcripts were checked for both the presence of transmembrane helices (TMHs) and for signal sequences, the first step towards identifying secreted effectors. A total of 12 candidate *D. citri* secreted effectors were found: five lack annotation or are otherwise *D. citri*-specific, and four were predicted to contain a TMH. Of the eight candidate salivary gland effector transcripts without TMHs, seven are highly up-regulated in *C*Las (+) adult *D. citri*, while one of the unknown transcripts is highly down-regulated in *C*Las (+) adult salivary glands. (Table S8). A recent paper by Wu et al [40] looked closely at salivary proteins and transcripts from CLas (-) *D. citri*, and of the eight possible effectors identified by this study, only the serine proteases were found in common.

#### Bacteriome

A key group of transcripts likely involved in communication between *D. citri* and its obligate endosymbionts housed in the bacteriome are the transporters, methyltransferases, acetyltransferases and the PiggyBac transposable elements, which together are represented in the top DE transcript list by 10 different transcripts. Three methyltransferases are all highly up-regulated in the *C*Las (+) adult bacteriome (methyltransferase family protein L2FC=7.48, phthiotriol dimycocerosates methyltransferase L2FC=5.48, and protein arginine N-methyltransferase L2FC=2.10) and one acetyltransferase is down-regulated (histone acetyltransferase catalytic subunit L2FC= -2.17). Five transcripts are annotated as “transporters” including three that are up-regulated in *C*Las (+), (cation-chloride cotransporter L2FC=3.07, cationic amino acid transporter L2FC=8.43, major facilitator transporter L2FC=5.59) and two that are down-regulated in *C*Las (+), (ABC transporter G family protein L2FC= -2.13 and organic solute transporter ostalpha protein L2FC= -2.31). Three ribosomal-related transcripts are up-regulated in the *C*Las (+) adult *D. citri* bacteriome (60S L26 with L2FC=4.03, 60S L37a with L2FC=3.25, and ribosomal protein L23 with L2FC=2.93). The full list of statistically significant (padj<0.05) bacteriome differentially expressed (L2FC>|2|) transcripts can be found in Table S5.

#### Head

The head dataset had relatively few reads sequenced and likewise, very few transcripts were statistically significantly DE. Of the 10 with padj<0.05 and L2FC>|2|, half (n=5) were associated with cell structure and signaling (including a vigilin gene with L2FC=-4.09, consistent with reports that *C*Las infection alters vector behavior [49]: a DNA-polymerase gene with L2FC=-3.42, a Rho-GTPase with L2FC=5.28, a neuromodulin gene with L2FC=5.27, and an insulin-like growth factor with L2FC=5.63). One transcript was associated with activation of autophagy (“Tumor protein p53-inducible nuclear protein 1” with L2FC=5.61), two with transport (an ATP synthase subunit gene with L2FC= -5.33, and one with an intracellular protein transport protein with L2FC= 3.41) and the final two were unknown (Dcitr10g06500.1.1 with L2FC= 5.56, and Dcitr05g06500.1.1 with L2FC= -4.26). Two overlaps between transcripts found in the salivary gland and head datasets included RNA-directed DNA polymerase which is highly down-regulated in *C*Las (+) adults in both datasets, as well as two ATP-synthase transcripts, one up-regulated in salivary glands (ATP synthase gamma chain L2FC=2.56), one down-regulated in heads (ATP synthase delta subunit L2FC= -5.33). The full list of statistically significant (padj<0.05) differentially expressed (L2FC>|2|) head transcripts can be found in Table S6.

### Genome improvement leads to quantifiable differences in RNAseq data interpretation

We hypothesized that due to improvements in the v3.0 *D. citri* genome, integrating across different datasets for visualization of tissue specific responses may have been successful in part due to improved transcript quantification. To test this hypothesis, the midgut dataset was used to compare RNAseq alignment and DE results between the v.1.1 and v.3.0 *D. citri* genome. The two versions of the *D. citri* genome resulted in different interpretations of the midgut transcriptomics results. The initial indication of differences between the two genome alignments was at the individual biological replicate level, where genome v3.0 has a 9% higher overall read alignment, as well as 3000 fewer *D. citri* transcripts found in each biological replicate, on average. The improved v3.0 genome changed the quantification of the midgut transcriptome. After differential expression, fewer statistically significant (adjusted p-value<0.05) differentially expressed transcripts (Log2FoldChange>|0.5|) were matched to genome v3.0 than genome v1.1. Percent alignment of cleaned reads was less than 100% in all biological replicates for both genomes (Table 2).

**Table 2:** Comparison of number of raw and trimmed reads from all biological replicates analyzed, as well as percent alignment, number of transcripts, and number of up and down regulated transcripts from both the v1.1 and v3.0 genome analysis of *D. citri C*Las (+) midguts.

| Raw read cleaning and filtering stats |  |  |  |  |  |  |
| --- | --- | --- | --- | --- | --- | --- |
| Midgut samples | #raw paired reads | #reads trimmed<sup>1</sup> | %aligned v1.1<sup>2</sup> | %aligned v3.0<sup>2</sup> | #transcripts v1.1<sup>3</sup> | #transcripts v3.0<sup>3</sup> |
| CLas(-)_1 | 27.85M | 273 | 64.89 | 73.82 | 17,170 | 13,814 |
| CLas(-)_2 | 28.26M | 234 | 68.13 | 77.12 | 15,284 | 12,481 |
| CLas(-)_3 | 26.05M | 246 | 66.12 | 74.29 | 17,566 | 14,142 |
| CLas(+)_1 | 26.89M | 76 | 64.04 | 73.23 | 16,339 | 13,281 |
| CLas(+)_2 | 27.15M | 210 | 62.16 | 71.82 | 16,834 | 13,641 |
| CLas(+)_3 | 22.41M | 117 | 64.48 | 74.77 | 16,476 | 13,230 |
| *D. citri* genome v1.1<sup>4</sup> |  |  | *D. citri* genome v3.0<sup>4</sup> |  |  |  |
| UP | DOWN | TOTAL | UP | DOWN | TOTAL |  |
| 272 | 341 | 20,792 | 176 | 303 | 12,704 |  |
| 1.30% | 1.64% | 100% | 1.38% | 2.38% | 100% |  |

<sup>1</sup>Trimming performed using Trimmomatic to remove adapters and low quality sequences.
<sup>2</sup>Alignment of cleaned reads to each genome performed using Hisat2. Quantities of single- and multi-aligning concordant reads were added together to calculate percent alignment.
<sup>3</sup>Transcripts were counted before differential expression and include only named, annotated Dcitr (v3.0) or XM (v1.1) IDs that have 1 or more counts. Not all transcripts are found in all biological replicates and not all are found in both CLas(+) and CLas(-).
<sup>4</sup>Differential expression performed via Ballgown and DESeq2. Transcripts in “TOTAL” column have at least 1 read aligning, while UP and DOWN regulated transcripts have adjusted p-value <0.05 and Log2FoldChange>0.5.

**Table 3:** Four statistically significant, differentially expressed genes from v3.0 midgut alignment were subject to BLAST to find their v1.1 genome equivalent gene IDs, and their total read counts, adjusted p-values, and Log2FoldChanges are compared.

| v3.0 Gene ID | v3.0 padj<sup>1</sup> | V3.0 Log2FC<sup>2</sup> | v1.1 Gene ID | v1.1 padj<sup>1</sup> | v1.1 Log2FC<sup>2</sup> |
| --- | --- | --- | --- | --- | --- |
| Dcitr10g01470.1.1 | 0.00 | -10.69 | LOC103515983 | 0.22 | -1.21 |
|  |  |  | LOC103515984 | 0.50 | -0.92 |
|  |  |  | LOC103518803* | 1.00 | 0.34 |
| Dcitr11g09870.1.1 | 0.01 | -0.511 | LOC103518620 | 0.14 | 1.60 |
| Dcitr13g03130.1.1 | 0.01 | -0.62 | LOC103509242 | 0.87 | -0.21 |
|  |  |  | LOC103509238 | 0.86 | -0.43 |
| Dcitr13g03190.1.1 | 0.01 | 0.51 | LOC103513428 | 0.72 | 0.66 |
|  |  |  | LOC103509249 | 0.84 | -0.44 |
|  |  |  | LOC103509235 | 0.51 | 0.56 |
|  |  |  | LOC113471714 | 0.55 | 0.53 |

<sup>1</sup>Adjusted p-values determined by DESeq2 using Benjamini-Hochberg adjustment of p-values.
<sup>2</sup>Log2FoldChange is calculated relative to healthy, so negative values show reduced expression in CLas (+) samples, while positive values show increased expression in CLas (+) samples.
\*Not enough read alignment counts for statistical analysis of differential expression.

Next, we hypothesized several possible ways the genome assembly could impact the interpretation of the transcriptome data (Figure 4A). The orange genome (representing version 1.1, Figure 4A) is shown in short fragments with variably sized gaps between the lengths. The reads from gene 1 (in blue) demonstrate multi-mapping to more than one genomic region, as well as non-alignment due to missing genomic sequence. The reads in green from gene 2 demonstrate that reads may align across a gap in the genome, and also that a dataset may not have reads to cover all the genome, or, alternatively the genomic sequence is such low quality that reads may not match to it perfectly enough to be counted. The corrected genome from v3.0 (pink) would be predicted to minimize these spurious mapping occurrences (Figure 4A, v3.0 genome in pink).

**Figure 4:**
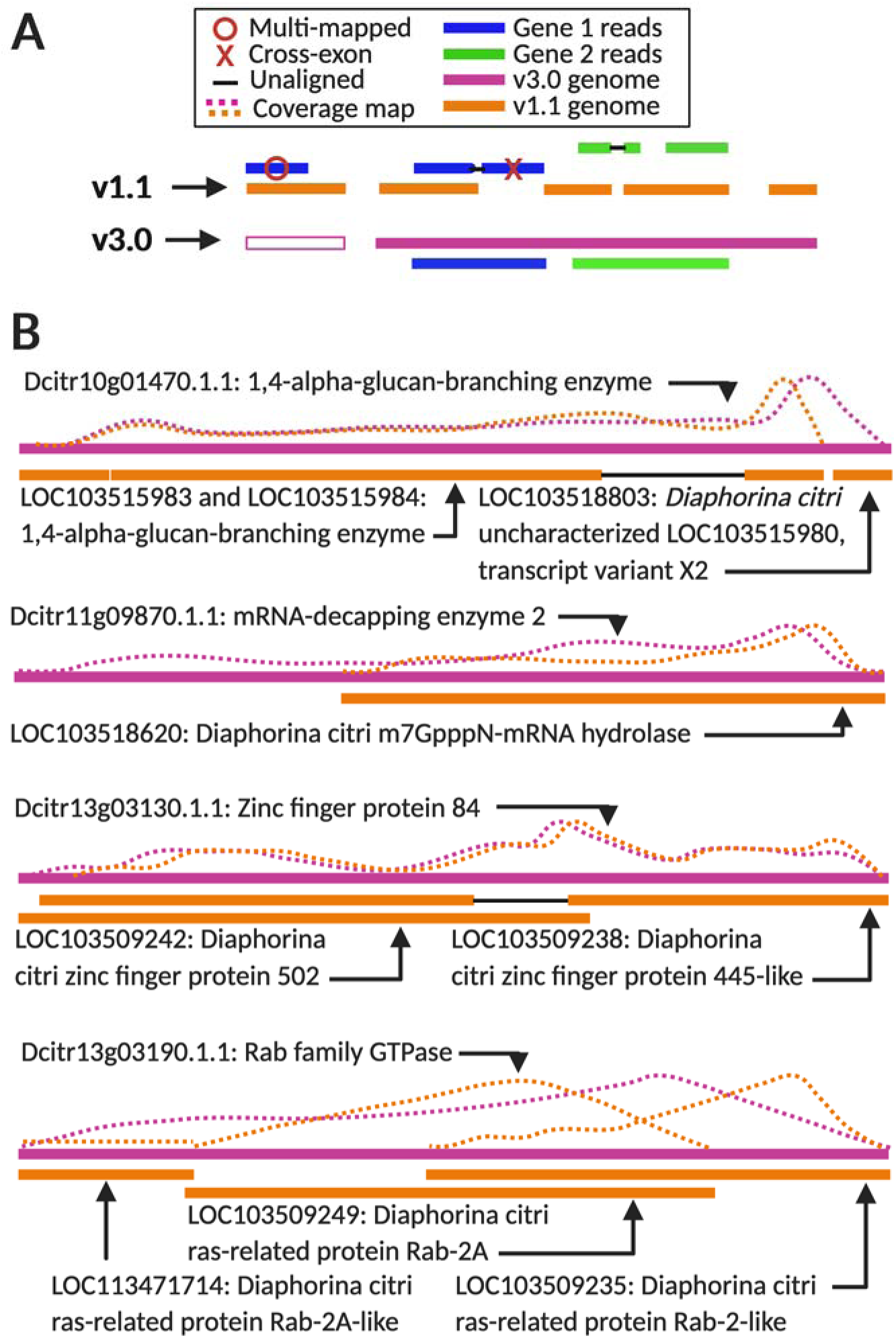
A) Predicted differences between the version Diaci_1.1 and v3.0 *D. citri* genomes. The genes in blue and green together demonstrate multi-mapping, non-alignment due to missing genomic sequence, alignment across a gap in the genome, and the genomic sequence is such low quality that reads may not match to it perfectly enough to be counted, while the updated genome represented in pink, fixes or reduces these issues. B) Four example transcripts showing differences in read alignment as a result of differences between the two genome versions. The pink line represents the newest genome v3.0 while orange represents the older genome, v1.1. Dotted lines demonstrate read alignment to the transcripts in the case of each genome.

To test whether these differences between genomes has a measurable effect on downstream expression analyses, we selected four random, differentially expressed transcripts (DE in the v3.0 analysis) for an in depth comparison (Figure 4B). As predicted, in all four cases, the new gene model was longer, and did not contain gaps. In contrast, the associated v1.1 gene models that matched to the full-length transcript were shorter, comprised of more fragments, included introns or gaps (Figure 4B), and were described as “PREDICTED” genes. We matched the read abundance profile over each transcript annotation to demonstrate differences in alignment frequency.

The transcript expression associated with each of the v1.1 LOC gene IDs which matched to the sequence from five differentially expressed transcripts from v3.0 (Fig. 4B) were assessed relative to v3.0 transcript expression. In all cases, the differential expression of the v1.1 transcripts in *C*Las-exposed relative to healthy was lower and less significant than the expression of the v3.0 transcripts.

## Discussion

The *D. citri* populations used to generate the samples in this study were all infected with *C*Las at different percentages, consistent with what has been reported in the literature for this pathogen and vector. Additionally, *C*Las reads were detected at high levels in the salivary gland and head samples, consistent with previous studies of the salivary glands using qPCR analysis [9, 11, 12]. The relative amount of *C*Las detected in the salivary gland data suggests that *C*Las is transcriptionally active, indicative of replication, though the lack of detection of similar numbers of *C*Las reads in the bacteriome and midgut does not preclude transcription, but that the levels may be below the limit of detection in these samples. Since sample RNA was poly-A enriched using oligos prior to making sequencing libraries, many of the *C*Las transcripts in samples are likely excluded, as poly-A tail enrichment biases samples towards eukaryotic mRNAs.

Detection of *C*Las reads in some tissues and not others leads us to revisit the nomenclature used to describe insects which are sampled from *C*Las-infected plants. Some studies, such as this one, designate insect samples as *C*Las (+) or *C*Las (-), or healthy or infected referring to the infection status of the tree used to rear the insect. Alternatively, some studies label insects (as opposed to the trees) as *C*Las-exposed or unexposed, the latter when sampled from healthy, *C*Las-negative trees. The use of exposed or unexposed is to account for the finding that not all insects acquire and/or become infected with *C*Las when reared on *C*Las-infected trees [10, 19, 20]. This transcriptomics study suggests that the exposed and unexposed designations are the most accurate because there is deeper complexity of *C*Las infection status in each insect at the level of the organ. In this study, salivary glands appear to have 10x more *C*Las reads than found in midguts and even more than in bacteriomes, suggesting salivary glands are truly “infected” and other organs, such as the bacteriome, remain “exposed”. Tissue specific gene regulation of vector-pathogen interactions was first described for poleroviruses that are transmitted by aphids [50]. Understanding the genetic basis, of both psyllid and bacteria, of the variation of infection in distinct organs in *C*Las (+) insects is an important new research frontier in this pathosystem. Improvements in RNAseq technology and *C*Las metagenomics will facilitate these types of studies.

It is more difficult to interpret whether organs are infected based on read count alone when read counts are barely above background, such as in the midguts. Kruse et al [15] reported that 82% (n=20, Cq<40) of the *C*Las (+) *D. citri* population which was harvested for their midguts were positive for *C*Las with an average qPCR Cq value of 31 across their four *C*Las (+) biological replicates. While 212 is not an especially large number of *C*Las reads post poly-A enrichment, when paired with the qPCR results, midguts, which have been shown to contain a visible slurry of *C*Las cells in previous work using microscopy [15, 17], may be referred to as “infected” by *C*Las, but at a lower level than the salivary glands. However, similar number of *C*Las reads were detected in the head samples from insects sampled from healthy (unexposed) trees as in the midguts. Finding a low level of reads aligning to *C*Las in healthy samples is not unexpected, and may be due to a few understandable reasons, such as a lack of enrichment of bacterial transcripts following poly-A enrichment for eukaryotic mRNAs, alignment errors, genome annotation errors, or homology of these reads to other psyllid-associated bacteria (the bacterial endosymbionts). *C*Las (-) psyllid colonies and citrus plants are reared in separate but identical environments to *C*Las (+) trees and insects. All materials are tested regularly and thoroughly for *C*Las using qPCR to rule out the possibility of *C*Las infection in these samples prior to experimentation.

Since significantly more reads aligned to *C*Las from the salivary gland dataset, these reads were mapped to known genes in the *C*Las-psy62 genome for annotation. All *C*Las transcripts had low read counts, most were unannotated, but two transcripts from the fig operon and one from the par operon were detected. The fig operon is part of the flagellum, and is involved in cell motility, cellular processes, chemotaxis, and overall mobility, [51] making it a potentially important gene when *C*Las interacts with its sub-cellular environment in the psyllid. Interestingly, a BLASTx analysis of the coding sequences of both the figB and figC transcripts produced homology to multiple Liberibacter species (figC %identity range of 72.93-84.33%, figB %identity range of 63.08-76.15%). The non-pathogenic *Liberibacter crescens* had the lowest identity (figC % identity = 67.67%, figB % identity = 56.92%) relative to the other Liberibacters, including “*Ca.* L. solanacearum”, *Ca.* L. americanus”, “*Ca.* L. africanus”, “*Ca.* L. europaeus” and “*Ca.* L. ctenarytainae”. These results support the hypothesis that the fig operon may be active in Liberibacter bacteria that are transmitted by pysllids.

The parB gene binds DNA and is part of the parABS system, which is known to play a role in bacterial chromosomal partitioning, cell cycle control and cell division, [52] and works by nicking supercoiled plasmid DNA at AT-rich regions and thus can act as a transcriptional regulator. While overall takeaways are limited due to the low number of reads aligned to this *C*Las gene, finding the par operon at relatively high expression when *C*Las is in the salivary glands of *D. citri* is consistent with the hypothesis of bacterial multiplication in this organ. [11] Due to the low number of *C*Las reads found in the other datasets, parB was not detected and thus relative expression could not be compared across tissues.

PCA analysis enabled a global visualization of the variation both within and across the datasets. The bi-axis separation between the four datasets as seen in Figure 2 can be partially explained by the average amount of *C*Las present (PC1) and by their sequencing (PC2). The head and bacteriome datasets were collected and multiplexed together but sequenced separately from the midgut and salivary gland datasets (which were also sequenced at different times). Head and salivary gland samples produced the highest number of reads aligning to *C*Las in the infected biological replicates, and bacteriome and midgut read counts were relatively low. Hosseinzadeh et al [21] quantified *C*Las titer in multiple organs of *D. citri* and found that bacteriomes contained a very low titer of *C*Las, with only the reproductive organs showing a lower titer. The bacteriome is highly specialized and designed to provide a place for replication of obligate bacteria. It is encased in a layer of psyllid cells (bacteriocytes), which could act as a barrier to *C*Las entry. Despite the lack of *C*Las in the bacteriome, it still had marked differences in the transcriptome between *C*Las(+) and *C*Las(-). For example, the Dcitr05g01800.11 transcript, has a log2FoldChange of 2.473, with a length 612 nucleotides, annotated as the “PiggyBac transposable element-derived protein 4”. It was significantly differentially expressed in the bacteriome dataset and not the other datasets. In the Diaci_v3.0 genome, this transcript is one of at least 11 PiggyBac-related genes found scattered across the genome (see Table S9). The PiggyBac (pB) transposon was first discovered 30 years ago in the cabbage looper, and now it is regularly used to transform insects, such as *Drosophila melanogaster*. PiggyBac is unique among transposases because of its specificity and seamless excision [53]. DNA between two sites with the specific sequence “TTAA” can be cleanly excised and the resulting DNA ends can perfectly match again without leaving a genomic footprint or synthesizing any new DNA. Similarly, the excised transposon can be re-integrated at any TTAA site in the genome. Due to the precision of pB, it is difficult to know exactly where Dcitr05g01800.11 originated – whether from the syncytial cytoplasmic cells, or the outer bacteriocytes. Considering what is known about pB and the bacteriome interactions with endosymbiotic bacteria, Dcitr05g01800.11 is a strong candidate for future studies of the bacteriome and using pB may open pathways for transgenesis in *D. citri*.

In addition to the bacteriome dataset, the infected midguts also recorded low *C*Las reads, (relative to salivary glands), which may be explained by general transcriptional inactivity of *C*Las during acquisition from the phloem. The relatively low replication rate in the midgut vs salivary glands may be an adaptive strategy to switch hosts from plant to insect to evade detection by the psyllid immune system [11, 54]. It is interesting that, although there were low levels of *C*Las reads in the midgut, the impact of *C*Las infection on the *D. citri* transcriptome was greatest in the midgut as compared to other tissues, which showed clear separation between *C*Las(+) and *C*Las(-) samples as a result of *C*Las infection. In adult insects, feeding on *C*Las- infected plants has been shown to induce drastic morphological changes to the psyllid nuclear architecture and apoptosis in the midgut epithelial cells [16, 17]. These data suggest that the infected plant sap, and not *C*Las directly, may be playing a role in modulating the midgut transcriptome response.

Given that the head samples were collected from a different cohort of insects than the salivary gland samples, and the salivary gland transcriptome is expected to be represented to some extent in the head transcriptome, the clustering of the head and salivary gland samples in PC1 was particularly encouraging and shows that transcriptome datasets collected in different experiments can be compared in the same analysis. The excised heads contained multiple organs which *C*Las-infected phloem or saliva pass through including the esophagus, foregut, mouthparts and salivary glands. *C*Las has been found in the brain [21], which is also present in head samples. Thus, the head may contain on average, a greater number of *C*Las bacteria than the other datasets as it contains more organs that *C*Las have been shown to inhabit. However, the head of the psyllid is a highly sclerotized part of the body. Sclerotization may have led to reduced yield when extracting nucleic acids due to reduced disruption efficiency and blockage of filters, two possibilities that may have led to the low yield – both of raw reads and alignment to the *D. citri* genome in these samples. Additionally, it has been shown that eye fluids of insects can contain PCR inhibitors that may interfere with library amplification and sequencing [55, 56].

The *D. citri* midgut has been a focal point of many studies. The initial draft genome fueled early studies to understand *C*Las-*D. citri* interactions and key insights into the vector-pathogen relationship have been derived from these studies. Researchers during the 2015-2020 time period used either Diaci_v1.0 or v2.0 for their analysis. We reanalyzed the raw, adult *D. citri* midgut data from both healthy (*C*Las-) and exposed (*C*Las+) samples originally published in 2016 by Kruse et al [15], using the v3.0 genome, just recently released [34]. As reported by Hosmani et al, [34], the version 1.1 genome contains 19.3Mb of gaps (Ns) and a large number (161,988) of short (<1Mb) scaffolds, making this assembly a useable but highly fragmented and incomplete picture of the genome. On the other hand, the version 3.0 genome is shorter, (473.9Mb) with less gaps (13.4Mb total Ns), and contains 13 chromosomal-length scaffolds (50.3Mb) and 1244 unplaced smaller scaffolds. The version 3.0 genome is also paired with, and improved by, curated and predicted gene and transcript annotations (totaling 19,049 genes and 21,345 transcripts) [33]. The improvements in overall read alignment rate of the midgut data to the v3.0 genome compared to the v1.1 genome suggests that, during alignment to the v1.1 genome, thousands of *D. citri* reads were completely left out of the analysis. The lower number of transcripts that matched to genome v3.0 is consistent with the increased scaffold length and gene model improvements.

The full-length transcript from the v3.0 analysis was searched against the v1.1 *D. citri* genome using BLAST (see methods). These analyses clearly show how quantification accuracy is improved with the full-length gene models, as all reads matching to a particular transcript are fully accounted for and used for differential expression analysis. Though each of these transcripts being analyzed is relatively short – comprising about 600-4000 nucleotides in length - the difference in read alignment frequency can be in the hundreds. We hypothesized that an improved genome sequence would change how transcriptomics results are interpreted. Analysis of a selected 4 transcripts showed this to be the case. In the v1.1 analysis, all 10 of these fragmented gene IDs and their associated transcripts would have been disregarded from the DE analysis because their adjusted p-values did not meet the significance threshold and the differential expression was nearly nonexistent (L2FC<|1|), and/or counts were too low and lacking in the biological replicates to be used. However, according to the v3.0 analysis, each of the four genes and their transcripts should be considered in downstream pathway analyses of effects of *C*Las exposure as they satisfied the adjusted p-value and log2FoldChange cutoffs. Thus, by quantifying how improved genome assemblies can lead to changes in differential expression, we present evidence to show that long read sequencing or other genome sequence improvement efforts are foundational for transcriptome-wide expression studies.

Improved genome quality did not; however, determine what proportion of differentially expressed transcripts were up or down regulated. The proportion of differentially expressed transcripts may be derived from the biology of the organisms or samples and in part the bioinformatic pipelines. Three studies look at the midgut of *D. citri* using transcriptomics: The analysis by Kruse et al. using v1.1 [15], this study using the Kruse et al data and the v3.0 genome, and a study by Yu et al. [41] using the v2.0 genome. The source of the midgut RNA is significantly different between the Yu et al. study and the Kruse et al study. Yu et al pooled midguts from *D. citri* adults raised on *Murraya exotica*, whereas Kruse et al. and thus, the current study, utilized insects raised on *Citrus medica*. Yu et al. also reported different *C*Las- infection rates among their individual insects pooled compared to Kruse et al. The relative proportions of transcripts that are up or down regulated in each of the three studies is not consistent, nor does the pattern become consistent with improved genome quality. In studies by Yu et al. and Kruse et al., there are more up regulated transcripts (499 and 965 respectively) than down regulated transcripts (279 and 850 respectively), while in this current study, the opposite is true (176 up and 303 down) (Table S3). The midgut analysis by Kruse et al. aligned RNA reads to the *D. citri* genome assembly v1.1 using the bioinformatic tools RSEM and bowtie2 for alignment, followed by edgeR and DESeq2 for differential expression calculations. The raw data from Kruse et al. was reanalyzed in the current study using the most recent versions of the bioinformatic tools Hisat2 (genome alignment), Stringtie (transcript assembly), Ballgown and DESeq2 (differential expression). These two bioinformatic pipelines differ in their alignment algorithms, statistical methods, and importantly their ability to identify false positive and negative differentially expressed transcripts.

## Potential Implications

*C*Las is uncultivable and methods to study *C*Las-*D. citri* interactions are challenging. Genome sequencing is a foundational tool for our exploration of the molecular interactions among *D. citri*, *C*Las, the bacterial endosymbionts and the citrus host. Our research showed that improved genome assemblies influences interpretation of transcriptomic data and that investigators have reason to re-analyze their previous *D. citri* transcriptomic data with the new genome release. The more accurate quantification provided by the Diaci_v3 genome may reduce the need to validate transcriptomic changes using reverse transcription (RT)-PCR. We urge arthropod genome communities and funding bodies to continue to invest funds on genome improvement projects such as i5k [57] and Ag100Pest [58]. These investments can help save expenditures elsewhere by reanalyzing previously generated and yielding higher confidence in the results after using a quality genome backbone. Additionally, single-cell RNAseq is the next frontier of understanding insect-pathogen interactions, especially for intracellular symbionts, at the highest resolution. Currently, single-cell RNAseq has been done on very few insects, but the list is expanding [59–62].

Still, a major roadblock is the functional annotation of the gene models. While automated pipelines for annotation exist at NCBI and elsewhere [63], these efforts are supplemented by manual annotation efforts [64–68] for *D. citri* and other arthropods [57]. Future work on understanding how the improved genome leads to improved quantification at the proteome level is also needed, and we hope such studies are inspired by the findings we present here.

## Methods

### *C*Las titer determination by qPCR

*D. citri C*Las-exposed and unexposed colonies were tested for the presence of *C*Las using qPCR by amplification of the 16S rDNA using TaqMan reagents. Individual, whole-body, adult psyllids (n=50 for the midgut colony, n=20 for the salivary gland colony, n=20 for the colony used to collect heads and bacteriomes) were collected from each colony. Total DNA was extracted from individual insects using the Qiagen DNeasy kit. DNA concentration was measured using a Nanodrop spectrophotometer. Each sample was standardized to 30 ng/ul so the Cq values from each dataset can be compared directly subjected. The *C*Las probe (5’-FAM-AGACGGGTG/ZEN/AGTAACGCG-3’) sequence and specific forward (5’-TCGAGCGCGTATGCAATACG-3’) and reverse (5’-GCGTTATCCCGTAGAAAAAGGTAG-3’) primers used are as published previously in Kruse et al. [15]. Unexposed colonies were tested monthly and *C*Las (+) colonies were tested at the time the insects were collected for dissection. Each qPCR plate contained positive and negative controls as well as a CLas 16S rDNA standard curve to allow for both absolute and relative *C*Las titer quantification, and every sample was run in triplicate. For our purposes, only Cq values were required to determine to whether individual samples were *C*Las (-/+) and to record the percent infection rate (how many out of 20 were *C*Las (+)) of the colony. A sample was considered *C*Las (+) if the Cq value was <40 (if there is only a single molecule in the reaction, with perfect primer efficiency, 37-40 cycles will be the cycle plateau). The Cq data from all 20 individuals, from all three colonies (bacteriomes and heads were collected from the same individuals and thus the same colony) was compiled and reported in **Figure S1**. Cq values from the CLas unexposed insects were undetected.

Once colonies were confirmed *C*Las (+) or *C*Las(-) by qPCR, hundreds of one to two week-old adult *D. citri* were collected and pooled to create biological replicates. Midgut samples included 250 guts pooled per biological replicate [n=3 replicates each *C*Las(+) and *C*Las(-)], salivary gland replicates each included 150 pooled extirpations [n=4 replicates each *C*Las(+) and *C*Las(-)], while 120 bacteriomes and heads were pooled for each replicate [n=5 replicates each *C*Las(+) and *C*Las(-)]. Salivary glands and midguts were pooled in TriZol while bacteriomes and heads were pooled in beta-mercaptoethanol and Qiagen RLT buffer. Samples were stored at - 80C until RNA extraction. The paired-end 150bp Illumina sequencing, raw data was uploaded to NCBI and is accessible to reviewers via BioProject accession # PRJNA385527, submission ID SUB10382129 and will be made available to the public upon publication.

### *In silico* quality control and cleaning of raw data to reduce confounding factors in analysis

Data analysis was conducted on servers hosted by the Computational Biology Center at the Boyce Thompson Institute. Data for all four datasets (bacteriome, head, salivary gland and midgut) were subjected to identical computational assessments and manipulations to eliminate variability caused by analysis methods. Total raw mRNA reads were first analyzed with FastQC [69] to gauge the presence of anomalies and adapters. Illumina Universal adapters that were present were removed by first interleaving/merging together forward and reverse reads into one large file. This file was then presented to AdapterRemover [70] using the Unix commands suggested in the manual for PE analysis. AdapterRemover output a file of interleaved paired-end reads that survived adapter removal. FastQC was run for the second time on this file to confirm adapter removal and check remaining read lengths and total remaining read quantity. This interleaved file was then used as input for SortMeRNA [71] which removes rRNA that survived the poly-A enrichment *in silico,* based on rRNA databases for bacteria, eukaryotes and archaea provided with the software program. Seed length was adjusted from default 18 down to 14 during rRNA database file indexing to be compatible with the minimum length reads in the current data set. SortMeRNA supplied two output types: 1) Those reads that mapped to rRNA (both forward and reverse reads had to map to be included), and 2) those where one or both of the paired end reads did not map to rRNA, such that the non-rRNA read pool contained some single strand sequences that aligned to rRNA. Separating out rRNA reduced over expression and bias of ribosomal gene expression in the datasets without totally removing rRNAs from the analysis. Low quality sequences (QC<20) were removed with Trimmomatic [72]. Paired reads where one or more are shorter than 17 nucleotides were then discarded. FastQC was run for the third time on these files to check their new read length distribution, read number and overall quality. A shell script was used to unmerge the forward and reverse reads for each sample file (reverse interleaving), creating a set of paired-end data files containing “cleaned reads” that could be used in the following steps.

### Read alignment to multiple genomes and differential transcript expression for each dataset

All four datasets comprised of cleaned, paired-end mRNA reads were aligned to both the v3.0 *D. citri* genome and the “*Candidatus* Liberibacter asiaticus” psy62 genome available on NCBI. The midgut dataset was additionally aligned to the v1.1 *D. citri* genome. The computational methods closely follow those published by Pertea et al., [73] and include the following: Each *D. citri* genome was indexed using HISAT2 (*hisat2-build*) [74]. Total cleaned reads were aligned to the indexed genome using *hisat2* and standard settings for PE data as described in the HISAT2 manual [74]. Specifically, options added to the base function included index memory mapping (--*mm*); setting the number of server threads to increase the speed of the alignment (-*p*); specifying output file names for both concordant alignments and non-concordant alignments (--*al-conc* and --*un-conc*, respectively); specifying which of the input files was forward or reverse (specified by “RF” showing -1 was reverse and -2 was forward); and tailored the output file organization for the possibility of downstream transcript assembly (--*dta*). Additionally, read alignment statistics were directed into a .stdout file for ease of future reference. Reads that aligned concordantly (collected in the *--al-conc* output file) were checked with FastQC and used in the next steps. Following alignment, the SAM files were converted to BAM to save space and then sorted by name using SAMtools [75]. Once sorted, reads were bundled into transcripts using Stringtie [76] based on their alignments and promptly re-aligned to the .GTF/.GFF file specific to each genome, containing information on all known genes for that genome. This process labeled each transcript with a specific Gene_ID, genomic location and information on introns/exons. Finally, using the number of transcripts that align to each gene, a count matrix was formed using Stringtie and ballgown [73] to allow downstream differential expression (DE) analysis between *C*Las (-) and *C*Las (+) replicates, paired with data visualization. Differential expression was performed in R (v3.3.3) using DESeq2 [77], following standard protocols (DE determined by setting *C*Las (-) as the denominator such that positive Log2FoldChange (L2FC) indicates greater expression in *C*Las (+) replicates and negative L2FC indicates reduced expression in *C*Las (+) replicates relative to *C*Las (-)). Because each dataset (except bacteriome and head) was collected and sequenced separately, normalizing the datasets to each other had too many experimental variables that were uncontrollable, so DE analysis for *C*Las (-/+) was performed separately for each dataset. DE results, like those of the qPCR Cq data, could be compared directly for transcripts within a dataset, while transcripts across datasets could be qualified, though no direct or quantitative comparison of expression could be made between datasets. Reads that aligned to *C*Las in the *C*Las (+) samples were counted and only certain transcripts of interest were analyzed further.

### Statistics and data visualization of results

A variety of statistical methods and data visualization tools were utilized. A principal components analysis (PCA) of all four datasets combined was performed in R (*prcomp* and *plot*) using a large transcript count matrix combining the transcript expression count matrices from the four datasets. The count data was minimally normalized by transcript counts per million and transcripts not present in both *C*Las (-) and *C*Las (+) replicates were removed. Individual PCA plots were also generated in R (*plotPCA* and *ggplot*) to show separation between *C*Las (-) and *C*Las (+) biological replicates, using the DESeq2 rlog-transformed transcript data for each dataset individually. Following PCA analysis, R was used to generate Volcano plots of the differentially expressed transcripts from each dataset individually, again using the DESeq2 rlog-transformed data. The L2FC of each DE transcript was plotted against the negative log of the adjusted p-value (-log(padj)) for the same transcript using *ggplot*.

The comparison of expression results from the midgut dataset when aligned to either v3.0 or v1.1 of the *D. citri* genome was started by choosing four transcripts present and expressed in both analyses. The two genomes presented different gene_IDs and genomic location coordinates which was problematic for direct comparison of changes in expression or even direct comparison of transcripts. The transcript sequence from Diaci_v3.0 was analyzed using BLASTx against the v1.1 genome to determine which v1.1 transcripts aligned to the v3.0 transcript and whether alignment was partial or full. To demonstrate differences in read distribution between the two genomes for each of the four transcripts and to show differential alignment frequencies, the v3.0 transcript sequences and associated v1.1 transcript sequences were used as a genome backbone and total cleaned reads were re-aligned to these sequences using HISAT2 to generate the .BAM files of read alignments for each transcript. Coverage maps were generated for each transcript using an R script (*BEDtools*) written by Dave Tang [78]. The general pattern of coverage from these coverage plots was duplicated in cartoon form on top of the respective transcript cartoon, to demonstrate the differences in read alignment location and frequency between the two *D. citri* genomes.

Potential secreted effectors were determined from the list of top DE transcripts of the salivary gland dataset by running two programs – SignalP-v5.0 [79] which accesses protein sequences for the presence of signal peptides, and Phobius [80] which detects both signal peptides and transmembrane helices (TMHs) from a protein sequence. Transcripts that putatively contained signal peptides but not TMHs were considered candidate salivary gland effector proteins.

## Additional Information Sections

Data Submission Information for Reviewers:

Title specific to data itself: The RNAseq data for the transcriptome analysis of multiple *D. citri* organs including the head, bacteriome and salivary gland.
Abstract: Paired-end 150bp mRNA-seq raw read files generated from pools of organs, which may include contaminants and/or other endosymbionts in addition to *Candidatus* Liberibacter asiaticus. Data is ideal for differential gene expression analysis.
Author list: Marina Mann, Surya Saha, Lukas A. Mueller, Michelle Heck Data types: poly-A enriched RNA, i.e transcriptome data
Organisms/Tissues of each data type: All data from *Diaphorina citri*, tissues include gut, bacteriome, salivary gland, and head plus thoracic segment 1 and antennae.
Estimate of dataset size: 120 G
File organization: Tar archive named “Dcitri_SG_BAC_HEAD_mRNA.tar” which contains three sub-archives in the following order, called “BACarchive.tar”, “HEADarchive.tar”, “SGarchive.tar”. Each sub archive contains gzipped fastq files for forward and reverse of every biological replicate.
Acknowledgments: Funding to generate samples and sequence them from Michelle Heck and Lukas Mueller, USDA-NIFA grants 2015-70016-23028 and 2020-70029-33199.

## Declarations

### List of abbreviations

All abbreviations have been defined in the manuscript.

### Consent for publication

Not Applicable.

### Competing interests

The author(s) declare that they have no competing interests

### Funding

This project was funded by NIFA Predoctoral Fellowship 2021-67011-35143 (MM) USDA-NIFA grants 2015-70016-23028 (MH and LM), 2020-70029-33199 (LM) and USDA ARS Project number 8062-22410-007-00-D (MH).

### Authors’ contributions

MM: Took part in, or led, all aspects including conceptualization, data curation, formal analysis, funding acquisition, investigation, methodology, validation, visualization and writing of original draft as well as review and editing.

SS: Funding acquisition, conceptualization, methodology, resources, writing - review and editing.

JMC: Visualization, writing – review and editing, data curation.

MP: Methodology, resources, software, writing – review and editing.

KM: Data curation, resources.

LC: Funding acquisition, project administration, resources, supervision.

WBH: Funding acquisition, project administration, resources, supervision, writing – review and editing.

LAM: Funding acquisition, project administration, methodology, conceptualization resources, supervision, writing – review and editing.

MH: Conceptualization, investigation, methodology, project administration, resources, supervision, validation, visualization, writing of original draft and reviews and edits.

## Acknowledgements

We thank Jaclyn Mahoney (Cornell University) for assistance with lab work, Dr. Angela Kruse (now at Vanderbilt University) for teaching Marina Mann how to extract RNA from psyllid organs while Dr. Kruse was a graduate student at Cornell, Tracy Bell and Hanna Mann from IRREC at Fort Pierce, FL for their assistance with excision of salivary glands. We are also grateful to Dr. Robert Krueger at the USDA ARS Citrus Germplasm Repository for providing the Heck lab with pathogen-free citrus seeds.

## Notes

https://www.ncbi.nlm.nih.gov/bioproject/PRJNA609978

### Competing Interest Statement

The authors have declared no competing interest.

