## Supplementary material for "Quantification of new and archived *Diaphorina citri* transcriptome data using a chromosomal length *D. citri* genome assembly reveals the vector’s tissue-specific transcriptional response to citrus greening disease": Figure S1

**Figure S1:** A histogram of CLas Cq values from individuals tested from each colony used to generate RNAseq data.

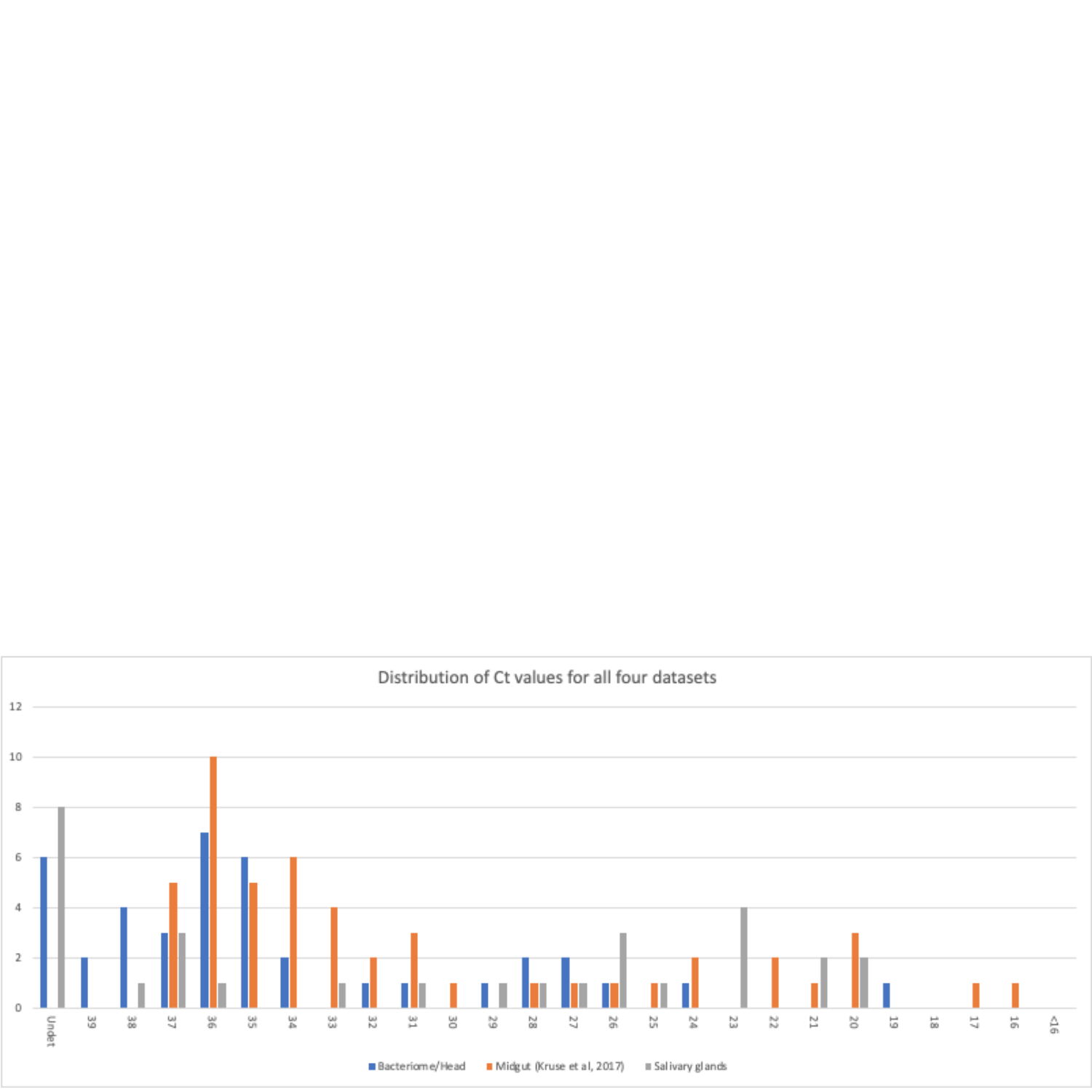
