## Supplementary material for "Quantification of new and archived *Diaphorina citri* transcriptome data using a chromosomal length *D. citri* genome assembly reveals the vector’s tissue-specific transcriptional response to citrus greening disease": Figure S2

**Figure S2**: PCA of all four datasets relative to each other, and distinguishing between healthy and *C*Las-exposed biological replicates.

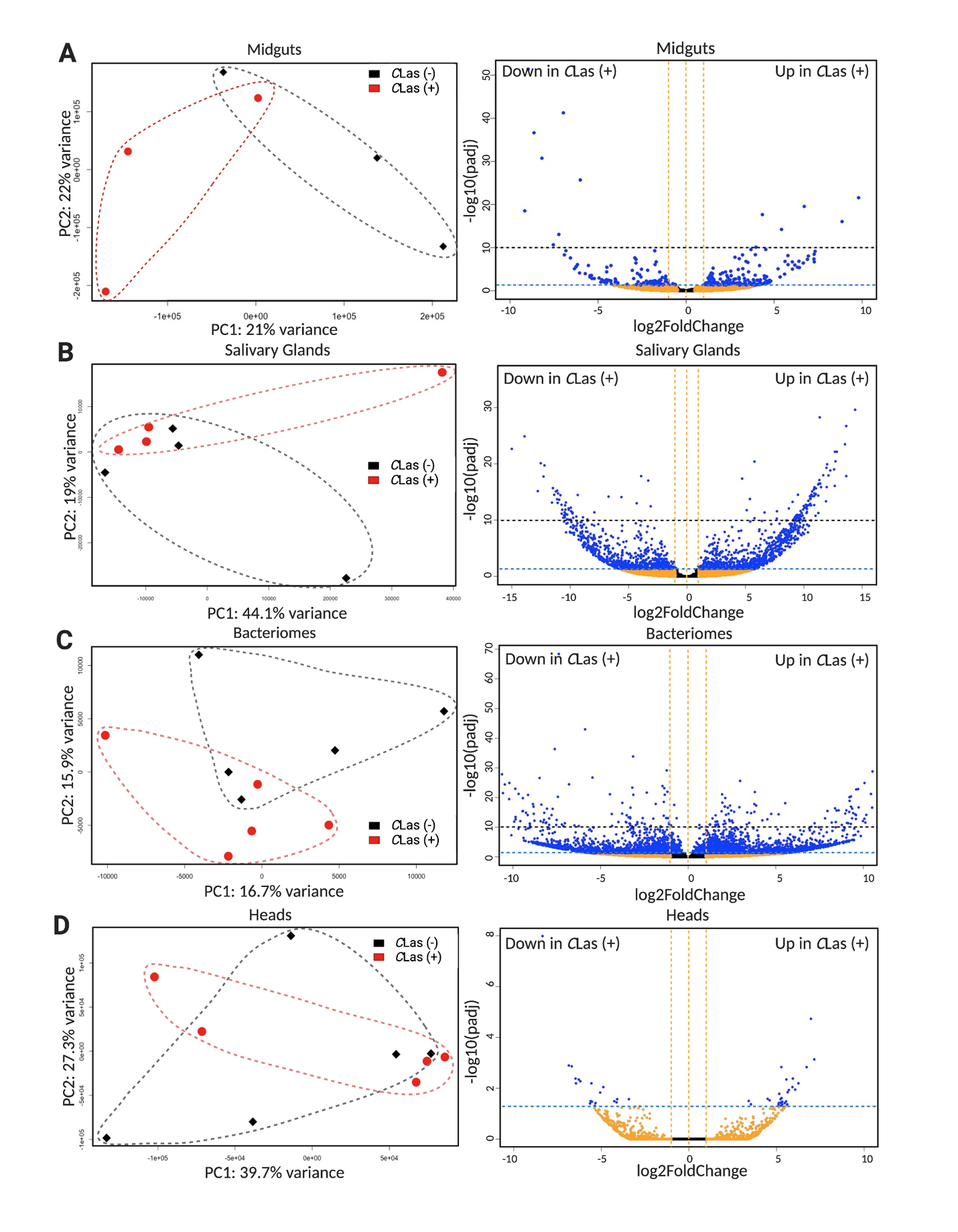
