## Supplementary material for "Quantification of new and archived *Diaphorina citri* transcriptome data using a chromosomal length *D. citri* genome assembly reveals the vector’s tissue-specific transcriptional response to citrus greening disease": Table S1

**Table S1**: Of the CLas transcripts identified from the salivary gland CLas (+) transcriptome dataset, only those CLas transcripts with reads present in three or more of the four biological replicates were considered (n=56). Among the top 10 with the most reads aligned, three transcripts stood out based on their annotation, figC, figB (two most abundant) and parB (9^th^ most abundant). All others are annotated as “unknown”. Reads were aligned to the CLas-psy62 genome from NCBI.

| **Order** | **Gene_ID** | **Total # reads aligned** | **Annotation** |
| --- | --- | --- | --- |
| 1 | gene269 | 290.00 | figB |
| 2 | gene268 | 176.00 | figC |
| 3 | gene911 | 166.00 | Unknown |
| 4 | gene702 | 148.00 | Unknown |
| 5 | gene1020 | 127.00 | Unknown |
| 6 | gene1056 | 126.00 | Unknown |
| 7 | gene788 | 87.00 | Unknown |
| 8 | gene543 | 77.00 | Unknown |
| 9 | gene416 | 71.00 | parB |
| 10 | gene993 | 80.00 | Unknown |
