## Supplementary material for "Quantification of new and archived *Diaphorina citri* transcriptome data using a chromosomal length *D. citri* genome assembly reveals the vector’s tissue-specific transcriptional response to citrus greening disease": Table S2

**Table S2:** Metadata on RNAseq datasets and alignments.

| **GUT** | **#**  **reads** | **%3.0** | **SG** | **#**  **reads** | **%3.0** | **BAC** | **#**  **reads** | **%3.0** | **HEA.** | **#**  **reads** | **%3.0** |
| --- | --- | --- | --- | --- | --- | --- | --- | --- | --- | --- | --- |
| *C*Las (-) 1 | 27.85 | 73.82 | *C*Las (-) 1 | 50.85 | 59.02 | *C*Las (-) 1 | 25.29 | 78.74 | *C*Las (-) 1 | 17.34 | 92.80 |
| *C*Las (-) 2 | 28.26 | 77.12 | *C*Las (-) 2 | 40.55 | 75.31 | *C*Las (-) 2 | 25.18 | 82.01 | *C*Las (-) 2 | 14.10 | 36.59 |
| *C*Las (-) 3 | 26.05 | 74.29 | *C*Las (-) 3 | 52.21 | 71.42 | *C*Las (-) 3 | 20.71 | 78.78 | *C*Las (-) 3 | 6.32 | 83.53 |
|  |  |  | *C*Las (-) 4 | 16.27 | 82.11 | *C*Las (-) 4 | 21.19 | 81.29 | *C*Las (-) 4 | 22.73 | 91.38 |
|  |  |  |  |  |  | *C*Las (-) 5 | 20.94 | 82.41 | *C*Las (-) 5 | 12.03 | 13.15 |
| *C*Las (+) 1 | 26.89 | 73.23 | *C*Las (+) 1 | 54.25 | 72.07 | *C*Las (+) 1 | 24.12 | 79.77 | *C*Las (+) 1 | 5.97 | 10.39 |
| *C*Las (+) 2 | 27.15 | 71.82 | *C*Las (+) 2 | 59.21 | 73.29 | *C*Las (+) 2 | 19.82 | 81.12 | *C*Las (+) 2 | 16.82 | 39.71 |
| *C*Las (+) 3 | 22.41 | 74.77 | *C*Las (+) 3 | 67.56 | 70.54 | *C*Las (+) 3 | 20.28 | 81.44 | *C*Las (+) 3 | 16.31 | 85.37 |
|  |  |  | *C*Las (+) 4 | 84.31 | 84.31 | *C*Las (+) 4 | 24.61 | 81.47 | *C*Las (+) 4 | 24.41 | 89.70 |
|  |  |  |  |  |  | *C*Las (+) 5 | 18.95 | 84.15 | *C*Las (+) 5 | 18.00 | 21.72 |
| Avg. | 26.43 | 74.17 |  | 44.98 | 73.51 |  | 22.11 | 81.12 |  | 15.40 | 56.43 |

GUT=Midgut dataset, SG=Salivary gland dataset, BAC=Bacteriome dataset, HEA.=Head dataset

#reads = The number millions of raw, paired-end reads.

%3.0 = The percent of paired-end reads that aligned to the version 3.0 *D. citri* genome.
