## Supplementary material for "Quantification of new and archived *Diaphorina citri* transcriptome data using a chromosomal length *D. citri* genome assembly reveals the vector’s tissue-specific transcriptional response to citrus greening disease": Table S3

**Table S3:** All transcripts from the midgut dataset that have differential expression log2FoldChange>|2| and adjusted p-value<0.05. Sorted by Log2FoldChange. Aligned to v3.0 of the *D. citri* genome.

**Transcript_ID	L2FC	padj	Annotation**

Dcitr10g01470.1.1	9.797	0.000	1,4-alpha-glucan-branching enzyme

Dcitr08g09790.1.1	8.859	0.000	Surfeit locus 4

Dcitr08g09800.1.1	7.327	0.000	SET and MYND domain-containing protein 4

Dcitr02g10600.1.1	7.283	0.000	Xanthine dehydrogenase

Dcitr08g03160.1.1	7.275	0.000	Murein tripeptide amidase MpaA

Dcitr03g01380.1.1	7.262	0.000	Alpha-mannosidase

Dcitr06g15560.1.2	6.946	0.000	centrosomal protein of 135 kDa, partial

Dcitr01g13610.1.1	6.891	0.000	Unknown protein

Dcitr03g07140.1.1	6.712	0.000	Acyl-CoA dehydrogenase

Dcitr08g02010.1.1	6.656	0.000	Open rectifier potassium channel protein 1

Dcitr02g11990.1.1	6.435	0.000	Gamma-glutamyl hydrolase

Dcitr04g05420.1.1	6.408	0.000	Unknown protein

Dcitr05g07620.1.1	6.329	0.000	Vinculin

Dcitr04g05810.1.1	6.173	0.000	28S ribosomal protein S18b, mitochondrial

Dcitr08g09260.1.2	6.039	0.000	Solute carrier family 41 member 2

Dcitr10g02030.1.1	6.034	0.000	UDP-glucuronosyltransferase

Dcitr12g06530.1.1	5.791	0.000	Unknown protein

Dcitr02g11330.1.1	5.783	0.000	DnaJ subfamily B member 2

Dcitr08g09460.1.3	5.618	0.000	Lipoma HMGIC fusion partner-like 2 protein

Dcitr09g06950.1.2	5.592	0.000	Nucleobase

Dcitr11g06490.1.1	5.420	0.000	zinc finger CCCH domain-containing protein 13-like

Dcitr01g06150.1.1	5.338	0.001	Mid1-interacting protein 1

Dcitr01g06150.1.2	5.338	0.001	Mid1-interacting protein 1

Dcitr03g09200.1.1	5.221	0.000	Metalloendopeptidase

Dcitr02g05500.1.2	4.803	0.006	neuropeptide CNMamide

Dcitr01g22250.1.1	4.762	0.008	RBR-type E3 ubiquitin transferase

Dcitr07g08080.1.1	4.744	0.004	Cysteine proteinase

Dcitr03g14920.1.5	4.724	0.005	GATA zinc finger domain-containing protein 15-like, partial

Dcitr01g12010.1.2	4.714	0.008	Pancreatic lipase-related protein 2

Dcitr08g09760.1.1	4.704	0.008	Insulin-like growth factor 2 mRNA-binding protein 3

Dcitr08g08680.1.1	4.701	0.013	Rho GTPase-activating protein 190

Dcitr06g06560.1.1	4.698	0.006	Intraflagellar transport particle protein 88

Dcitr03g17770.1.1	4.629	0.011	Unknown protein

Dcitr02g09070.1.1	4.625	0.016	Carboxypeptidase D

Dcitr02g09710.1.1	4.571	0.001	E3 ubiquitin ligase PARAQUAT TOLERANCE 3-like

Dcitr11g08160.1.1	4.568	0.005	Unknown protein

Dcitr10g01310.1.1	4.557	0.013	Ankyrin repeat protein

Dcitr02g07600.1.1	4.524	0.000	Enolase

Dcitr01g12820.1.1	4.478	0.013	ran-binding protein 3-like

Dcitr04g04620.1.1	4.452	0.001	Unknown protein

Dcitr02g07110.1.1	4.442	0.000	Phosphate transporter

Dcitr04g09180.1.3	4.411	0.023	selection and upkeep of intraepithelial T-cells protein 1-like isoform X2

Dcitr03g11060.1.1	4.333	0.000	ATP-dependent RNA helicase

Dcitr07g08110.1.1	4.323	0.026	Cathepsin

Dcitr01g13440.1.1	4.281	0.026	Unknown protein

Dcitr11g03350.1.1	4.240	0.033	Alpha-tocopherol transfer protein-like

Dcitr00g02160.1.1	4.224	0.003	CG12123-RA

Dcitr04g06690.1.1	4.205	0.017	Long-chain acyl-CoA synthetase

Dcitr01g01630.1.2	4.190	0.008	Bestrophin homolog

Dcitr01g05670.1.1	4.187	0.036	Lachesin

Dcitr01g18090.1.1	4.183	0.033	Unknown protein

Dcitr02g09760.1.1	4.176	0.033	Unknown protein

Dcitr11g01690.1.1	4.172	0.000	Unknown protein

Dcitr07g07340.1.1	4.162	0.033	circadian locomoter output cycles protein kaput

Dcitr04g09730.1.1	4.107	0.036	Odorant-binding protein 18

Dcitr01g09430.1.1	4.059	0.004	Protein prgI

Dcitr07g10360.1.1	4.029	0.035	Ankyrin repeat and fibronectin type-III domain-containing protein 1

Dcitr04g11360.1.2	4.028	0.004	Proton-coupled amino acid transporter 4

Dcitr02g03230.1.1	4.027	0.004	CLIPC3 -Serine Protease Snake-Like.

Dcitr07g06300.1.1	3.998	0.041	General secretion pathway protein C

Dcitr11g04950.1.2	3.967	0.000	Dynamin-like protein, mitochondrial

Dcitr00g04570.1.3	3.934	0.000	Zinc

Dcitr03g09760.1.1	3.922	0.000	UDP-glucuronosyltransferase

Dcitr03g11640.1.1	3.880	0.049	2,3-bisphosphoglycerate-dependent phosphoglycerate mutase

Dcitr00g01710.1.1	3.858	0.012	Major

Dcitr10g07760.1.1	3.822	0.003	Unknown protein

Dcitr10g02970.1.1	3.776	0.024	Unknown protein

Dcitr04g11510.1.1	3.774	0.029	Unknown protein

Dcitr05g12140.1.1	3.750	0.040	Vacuolar protein sorting-associated protein 37A

Dcitr05g13670.1.1	3.685	0.000	Dosage compensation regulator

Dcitr02g13200.1.1	3.647	0.030	Unknown protein

Dcitr06g07920.1.1	3.633	0.012	Unknown protein

Dcitr12g03760.1.1	3.621	0.017	Cathepsin L1-like Prot 4

Dcitr01g08080.1.2	3.614	0.014	Hydroxymethylglutaryl-CoA lyase

Dcitr13g01820.1.1	3.602	0.018	SITE-1 protease

Dcitr04g06130.1.1	3.547	0.000	LYR motif-containing protein 7

Dcitr12g03060.1.1	3.509	0.004	Unknown protein

Dcitr03g07590.1.1	3.508	0.000	serine/threonine-protein phosphatase 4 regulatory subunit 3

Dcitr10g09310.1.1	3.431	0.047	Ubiquitin-conjugating enzyme

Dcitr02g14920.1.1	3.400	0.000	Adenine phosphoribosyltransferase

Dcitr05g04040.1.1	3.362	0.015	Tumor protein p53-inducible nuclear protein 1

Dcitr02g12750.1.1	3.333	0.033	Mediator of RNA polymerase II transcription subunit 9

Dcitr03g05350.1.1	3.331	0.006	Protein bric-a-brac 2

Dcitr13g02990.1.1	3.310	0.001	Unknown protein

Dcitr07g10650.1.1	3.301	0.027	transcription factor SPT20 homolog, partial

Dcitr08g11710.1.2	3.241	0.000	3-hydroxy-3-methylglutaryl coenzyme A synthase

Dcitr00g05130.1.1	3.220	0.000	Unknown

Dcitr06g09810.1.1	3.208	0.036	WD repeat-containing protein on Y chromosome

Dcitr08g05230.1.1	3.202	0.000	Arylsulfatase B

Dcitr03g07160.1.1	3.173	0.033	La

Dcitr03g03770.1.1	3.062	0.021	GTP-binding protein 1

Dcitr02g20050.1.1	3.062	0.015	Tetratricopeptide repeat (TPR)-like superfamily protein

Dcitr07g03880.1.1	3.046	0.049	Unknown protein

Dcitr03g15760.1.1	3.022	0.014	Trypsin-like serine protease

Dcitr02g11140.1.1	3.017	0.000	keratin-associated protein 19-2-like isoform X1

Dcitr10g07700.1.1	3.016	0.002	Unknown protein

Dcitr08g07480.1.3	3.000	0.001	villin-2-like

Dcitr13g01890.1.1	2.963	0.000	Adenylate kinase isoenzyme 6 homolog

Dcitr10g07150.1.1	2.902	0.017	Heat Shock Protein 70 A1

Dcitr02g07960.1.1	2.806	0.002	Aquaporin AQPAe.a-like

Dcitr10g06510.1.1	2.774	0.000	Heat Shock Protein 70 B

Dcitr01g22530.1.1	2.746	0.018	Unknown protein

Dcitr11g01900.1.1	2.713	0.002	Alpha-(1,6)-fucosyltransferase

Dcitr08g03010.1.1	2.676	0.010	Peritrophin-1

Dcitr04g17010.1.1	2.659	0.012	Histone H3

Dcitr06g02580.1.1	2.599	0.019	Cobalt-zinc-cadmium efflux system protein

Dcitr08g07480.1.4	2.557	0.000	villin-2-like

Dcitr13g03230.1.1	2.535	0.000	cytochrome B561%2C amino-terminal protein Chr1:2469528-2472518 REVERSE LENGTH%3D693 201606

Dcitr13g01840.1.1	2.513	0.015	Adenylate kinase isoenzyme 6 homolog

Dcitr10g01270.1.1	2.489	0.022	4-aminobutyrate aminotransferase

Dcitr03g01700.1.1	2.488	0.003	72 kDa type IV collagenase

Dcitr05g09020.1.1	2.478	0.031	DNA-directed RNA polymerases I/II/III subunit

Dcitr00g10110.1.1	2.449	0.004	zinc

Dcitr11g01700.1.1	2.380	0.028	trypsin I-P1-like, partial

Dcitr05g10240.1.1	2.369	0.001	Phosphoenolpyruvate carboxykinase

Dcitr10g07040.1.1	2.343	0.018	Heat shock protein 70

Dcitr06g09500.1.1	2.316	0.000	Neurotrypsin

Dcitr04g14800.1.1	2.294	0.005	Major facilitator, sugar transporter-like, Major facilitator superfamily domain protein

Dcitr01g12250.1.1	2.277	0.044	Calmodulin

Dcitr01g22110.1.1	2.263	0.026	Mite allergen Der p 7

Dcitr01g13510.1.1	2.247	0.035	WD repeat-containing protein 19

Dcitr01g03180.1.1	2.240	0.000	Odorant binding protein

Dcitr04g17030.1.1	2.240	0.049	Histone H2B

Dcitr01g09440.1.1	2.212	0.040	WD40 repeat protein

Dcitr01g16420.1.1	2.143	0.040	Carboxypeptidase B

Dcitr05g13810.1.1	2.137	0.012	Dynein regulatory complex subunit 7

Dcitr01g06830.1.1	2.111	0.013	Protein with WD-40 repeat domain

Dcitr03g19110.1.1	2.075	0.009	Unknown protein

Dcitr03g11520.1.1	2.073	0.000	Cystatin

Dcitr08g10620.1.1	-2.030	0.007	Syntaxin-41

Dcitr08g09060.1.1	-2.059	0.014	Pleckstrin homology domain-containing family M member 2

Dcitr01g18410.1.1	-2.133	0.011	Cullin-associated NEDD8-dissociated protein 1

Dcitr03g13820.1.1	-2.185	0.000	PRELI/MSF1 domain-containing protein

Dcitr07g07760.1.1	-2.216	0.026	Doubletime

Dcitr03g15900.1.1	-2.257	0.006	Serine/threonine-protein phosphatase 4 regulatory subunit 3

Dcitr06g05800.1.1	-2.413	0.014	DNA-directed RNA polymerase III subunit rpc-3

Dcitr02g11330.1.4	-2.534	0.000	DnaJ subfamily B member 2

Dcitr10g09050.1.2	-2.563	0.001	Acyl carrier protein

Dcitr10g01500.1.1	-2.568	0.032	Zinc finger MYM-type protein 1

Dcitr03g19770.1.1	-2.569	0.033	Unknown protein

Dcitr03g06820.1.1	-2.581	0.049	Suppressor of fused

Dcitr08g02060.1.1	-2.640	0.000	RING/U-box superfamily protein

Dcitr03g07620.1.1	-2.729	0.008	DNA repair/transcription protein mms19

Dcitr13g03440.1.1	-2.772	0.000	Long chain acyl-coa synthetase

Dcitr01g10860.1.1	-2.826	0.037	Ubiquinol-cytochrome c reductase complex chaperone CBP3

Dcitr05g06210.1.2	-2.847	0.000	Unknown protein

Dcitr11g01280.1.1	-2.851	0.000	Zinc finger protein 879

Dcitr04g15470.1.1	-2.911	0.000	Tensin

Dcitr10g08880.1.1	-2.981	0.002	cold shock domain protein 1 Chr4:17043443-17044342 REVERSE LENGTH%3D299 201606

Dcitr11g09700.1.1	-3.057	0.000	Collagen alpha 2(I) chain

Dcitr05g05120.1.1	-3.347	0.033	Unknown protein

Dcitr00g01590.1.1	-3.878	0.015	Collagen

Dcitr01g05780.1.1	-3.886	0.015	Thymidylate kinase

Dcitr07g08900.1.1	-4.005	0.004	Phosphomannomutase

Dcitr08g01460.1.1	-4.024	0.002	HSP20-like chaperone

Dcitr06g14110.1.1	-4.026	0.010	Transmembrane protein 165

Dcitr03g13500.1.1	-4.245	0.028	MAM domain-containing protein 2

Dcitr00g06180.1.2	-4.265	0.000	Broad-complex

Dcitr08g12680.1.2	-4.271	0.005	Mediator of RNA polymerase II transcription subunit 14

Dcitr02g15740.1.4	-4.292	0.006	Rap guanine nucleotide exchange factor 4

Dcitr03g18670.1.1	-4.383	0.004	CBS domain-containing protein / transporter associated domain-containing protein

Dcitr01g04810.1.1	-4.393	0.004	26S proteasome regulatory subunit

Dcitr04g12470.1.1	-4.481	0.004	39S ribosomal protein L10, mitochondrial

Dcitr11g06930.1.1	-4.505	0.000	Thymidylate kinase

Dcitr01g11490.1.1	-4.512	0.000	28S ribosomal protein S24, mitochondrial

Dcitr05g17080.1.1	-4.516	0.010	Long chain base biosynthesis protein 1-like

Dcitr04g11590.1.1	-4.522	0.014	Suppressor of hairless protein

Dcitr05g15560.1.1	-4.525	0.003	Proton-coupled amino acid transporter 4

Dcitr10g08700.1.1	-4.613	0.012	CAP-Gly domain-containing linker protein 1

Dcitr01g19850.1.1	-4.662	0.009	Larval cuticle protein 16/17

Dcitr05g03690.1.1	-4.683	0.005	histidine-rich glycoprotein

Dcitr10g08260.1.1	-4.784	0.003	Sin3 histone deacetylase corepressor complex component SDS3

Dcitr03g08640.1.1	-4.842	0.010	GM25696

Dcitr06g04860.1.1	-4.944	0.001	LOW QUALITY PROTEIN: myb-like protein P

Dcitr03g12480.1.4	-5.217	0.001	S-adenosyl-L-methionine-dependent methyltransferases superfamily protein Chr1:5687994-5690395 FORWARD LENGTH%3D382 201606

Dcitr11g05020.1.2	-5.346	0.001	Trafficking protein particle complex subunit

Dcitr02g13040.1.1	-5.377	0.000	CD5 antigen-like

Dcitr06g14230.1.1	-5.452	0.000	Forkhead box protein O

Dcitr00g08930.1.1	-5.464	0.001	Serine

Dcitr00g04570.1.4	-5.549	0.000	Zinc

Dcitr02g14470.1.1	-5.729	0.000	Cell adhesion molecule 3

Dcitr05g04450.1.1	-6.012	0.000	Nuclear transport factor 2

Dcitr11g01720.1.2	-6.054	0.000	Alpha-tocopherol transfer protein-like

Dcitr01g10250.1.1	-6.174	0.000	Zinc finger protein 84

Dcitr04g09190.1.1	-6.248	0.000	jerky protein homolog-like, partial

Dcitr02g07060.1.1	-6.340	0.000	Dorsal-ventral patterning protein Sog

Dcitr10g01320.1.3	-6.570	0.000	Unknown protein

Dcitr10g09890.1.1	-6.828	0.000	Thioredoxin-mitochondrial 1

Dcitr04g15090.1.1	-6.913	0.000	NADH-quinone oxidoreductase subunit B

Dcitr00g04130.1.1	-6.971	0.000	Origin

Dcitr02g07360.1.4	-7.227	0.000	Adenylate kinase

Dcitr06g01400.1.1	-7.544	0.000	Unknown protein

Dcitr01g13900.1.1	-8.191	0.000	Mitochondrial pyruvate carrier

Dcitr01g14790.1.1	-8.641	0.000	Calreticulin

Dcitr02g11330.1.2	-8.972	0.000	DnaJ subfamily B member 2

Dcitr04g16360.1.1	-9.165	0.000	EH domain-binding protein 1
