## Supplementary material for "Quantification of new and archived *Diaphorina citri* transcriptome data using a chromosomal length *D. citri* genome assembly reveals the vector’s tissue-specific transcriptional response to citrus greening disease": Table S4

**Table S4:** All transcripts from the salivary gland dataset that have differential expression log2FoldChange>|2| and adjusted p-value<0.05. Sorted by Log2FoldChange.

**Transcript_ID	L2FC	padj	Annotation**

Dcitr04g09440.1.1	11.361	0.000	NADH-ubiquinone oxidoreductase

Dcitr04g13690.1.1	10.523	0.000	40S ribosomal protein S28

Dcitr10g10180.1.1	10.129	0.000	40S ribosomal protein S15a

Dcitr08g07040.1.1	9.306	0.000	Phosphate acyltransferase

Dcitr01g05060.1.1	8.412	0.000	Acyl-CoA thioesterase

Dcitr12g05490.1.1	8.155	0.000	Gamma-glutamylcyclotransferase

Dcitr01g09640.1.1	8.048	0.000	Alpha-tocopherol transfer protein-like protein

Dcitr10g02950.1.1	7.829	0.000	Signal peptidase I

Dcitr13g03480.1.1	7.406	0.000	Ubiquitin conjugating enzyme

Dcitr03g19180.1.1	7.209	0.000	CAAX prenyl protease 1 (Peptidase family M48)

Dcitr03g16830.1.3	7.171	0.000	basic salivary proline-rich protein 3-like

Dcitr05g11190.1.1	7.048	0.000	Methylthioribulose-1-phosphate dehydratase

Dcitr11g07970.1.2	6.765	0.000	Calcium-binding EF-hand

Dcitr13g07090.1.3	6.693	0.000	Farnesyl pyrophosphate synthase

Dcitr05g07830.1.1	6.622	0.000	LIM homeobox transcription factor 1-beta

Dcitr07g02170.1.1	6.442	0.002	Mitochondrial carrier

Dcitr01g11050.1.2	6.436	0.000	Soluble calcium-activated nucleotidase 1

Dcitr04g16010.1.1	6.260	0.002	Unknown protein

Dcitr11g09190.1.4	6.220	0.000	Geranylgeranyl transferase type-1 subunit beta

Dcitr13g03250.1.1	6.199	0.002	Leucine--tRNA ligase

Dcitr13g04800.1.1	6.048	0.000	Tethering factor for nuclear proteasome STS1

Dcitr08g04630.1.1	5.991	0.000	UDP-N-acetylglucosamine pyrophosphorylase 1

Dcitr02g12920.1.2	5.959	0.007	ABC transporter C family

Dcitr07g06610.1.1	5.910	0.000	Meteorin-like protein

Dcitr03g05340.1.1	5.892	0.001	tctex1 domain-containing protein 1-like

Dcitr05g14340.1.1	5.883	0.002	Flavin-containing monooxygenase

Dcitr03g15750.1.1	5.861	0.011	Serine protease

Dcitr01g11640.1.1	5.787	0.021	Serine/threonine-protein kinase 16

Dcitr13g01840.1.1	5.782	0.002	Adenylate kinase isoenzyme 6 homolog

Dcitr03g04260.1.1	5.774	0.000	Ribosomal protein

Dcitr06g03150.1.1	5.737	0.004	Phospholipid-transporting ATPase

Dcitr06g05570.1.1	5.727	0.002	NADH dehydrogenase [ubiquinone] 1 α subcomplex subunit 7

Dcitr01g21740.1.1	5.624	0.021	Thioredoxin domain-containing protein 17

Dcitr03g04400.1.1	5.535	0.001	60S ribosomal protein L4

Dcitr01g04160.1.1	5.462	0.000	Unknown protein

Dcitr02g18560.1.1	5.450	0.007	Unknown protein

Dcitr11g05250.1.1	5.428	0.003	Elongation factor 2

Dcitr05g04310.1.1	5.399	0.003	AGAP013432-PA

Dcitr06g05040.1.1	5.372	0.012	Unknown protein

Dcitr12g02110.1.1	5.359	0.029	Very-long-chain 3-oxoacyl-CoA reductase 1

Dcitr00g13000.1.1	5.234	0.006	ABC

Dcitr02g16560.1.1	5.193	0.015	TBC1 domain family member 31-like isoform X2

Dcitr10g09840.1.1	5.168	0.043	testis specific serine/threonine protein kinase 3

Dcitr01g13970.1.1	5.168	0.005	Elongation factor 1-beta

Dcitr01g10480.1.1	5.140	0.026	33 kDa inner dynein arm light chain, axonemal

Dcitr04g11380.1.1	5.059	0.040	Unknown protein

Dcitr03g02880.1.1	5.055	0.045	spermatogenesis-associated protein 7-like isoform X1

Dcitr01g05220.1.1	5.048	0.009	UPF0691 protein C9orf116

Dcitr03g04980.1.1	5.042	0.024	zinc finger protein 436-like isoform X1

Dcitr01g20800.1.1	5.006	0.029	Pleckstrin homology domain-containing family M member 1

Dcitr09g07010.1.1	4.986	0.011	Alpha-Crystallin

Dcitr10g02810.1.1	4.963	0.038	Unknown protein

Dcitr00g14590.1.1	4.908	0.021	Transposon

Dcitr11g08720.1.1	4.881	0.045	Protein yellow

Dcitr01g15810.1.1	4.856	0.025	RNA helicase

Dcitr05g09660.1.1	4.854	0.030	Unknown protein

Dcitr04g10840.1.1	4.841	0.015	Protein FAM210A

Dcitr13g01100.1.1	4.811	0.001	Snurportin-1

Dcitr03g11970.1.1	4.763	0.030	DNA repair RAD51-like protein

Dcitr10g07900.1.1	4.757	0.022	Macrophage erythroblast attacher

Dcitr08g10720.1.1	4.697	0.038	Homeobox protein Nkx-2.2a

Dcitr02g04500.1.1	4.694	0.030	Phosphoribosylamine--glycine ligase

Dcitr06g04060.1.1	4.693	0.031	Ubiquitin-ligase E3

Dcitr01g07660.1.1	4.670	0.028	Unknown protein

Dcitr05g14800.1.1	4.595	0.010	Elongation factor 4

Dcitr10g10090.1.1	4.579	0.038	Unknown protein

Dcitr01g16880.1.1	4.567	0.042	WD40 repeat

Dcitr02g04530.1.1	4.464	0.006	Biogenesis of lysosome-related organelles complex 1 subunit 1

Dcitr03g13410.1.1	4.445	0.021	Polyadenylate-binding protein-interacting protein 2B

Dcitr01g07920.1.1	4.398	0.024	Unknown protein

Dcitr06g11690.1.1	4.324	0.002	Peroxiredoxin

Dcitr03g06640.1.1	4.312	0.028	Zinc finger CCHC domain-containing protein 4

Dcitr02g13860.1.1	4.277	0.020	RNA-directed DNA polymerase from mobile element jockey

Dcitr08g03350.1.1	4.260	0.030	Ecdysone-induced protein 74EF isoform B

Dcitr04g11610.1.2	4.038	0.027	MFS-type transporter C6orf192

Dcitr07g08050.1.1	3.944	0.046	Transmembrane emp24 domain-containing protein 7

Dcitr10g06970.1.2	3.701	0.006	U3 small nucleolar RNA-associated protein 6

Dcitr03g15730.1.1	3.646	0.010	CLIPB - Serine Protease 1

Dcitr07g02410.1.1	3.265	0.032	Chloride channel protein

Dcitr05g16750.1.1	3.236	0.006	Pro-Pol polyprotein

Dcitr04g06660.1.1	3.182	0.035	NADPH-dependent diflavin oxidoreductase 1

Dcitr05g16000.1.2	3.180	0.049	Fatty-acid amide hydrolase 2

Dcitr13g04790.1.1	2.952	0.020	Lipid phosphate phosphatase

Dcitr02g08670.1.1	2.904	0.019	Lamin-B1

Dcitr04g09870.1.1	2.678	0.008	RNA polymerase II-associated protein 1

Dcitr10g07230.1.2	2.608	0.009	Ankyrin repeat and MYND domain-containing protein 2

Dcitr07g02010.1.1	2.587	0.008	Guanine deaminase

Dcitr07g02260.1.1	2.536	0.006	Nuclear autoantigenic sperm protein

Dcitr12g07070.1.1	2.530	0.013	oxidative stress-responsive serine-rich protein 1

Dcitr12g05190.1.1	2.256	0.015	ATP synthase gamma chain

Dcitr04g11810.1.1	2.218	0.032	Transposon Ty3-I Gag-Pol polyprotein

Dcitr04g02090.1.1	-2.522	0.025	Unknown protein

Dcitr07g03910.1.1	-2.647	0.001	Unknown protein

Dcitr11g01500.1.1	-3.290	0.000	RNA-directed DNA polymerase from mobile element jockey

Dcitr10g10160.1.1	-4.359	0.024	Protein sidekick

Dcitr03g13400.1.2	-4.409	0.000	UDP-glucuronosyltransferase

Dcitr09g07790.1.1	-4.641	0.045	conserved

Dcitr01g09640.1.2	-5.198	0.037	Alpha-tocopherol transfer protein-like protein

Dcitr02g02390.1.2	-5.323	0.010	E3 SUMO-protein ligase PIAS3

Dcitr08g05310.1.1	-5.772	0.043	5-aminolevulinate synthase

Dcitr02g02120.1.1	-5.908	0.000	Unknown protein

Dcitr05g07530.1.1	-6.262	0.017	U2 small nuclear ribonucleoprotein B

Dcitr06g05250.1.1	-7.423	0.000	Zinc finger protein 417

Dcitr03g04630.1.2	-7.934	0.000	NAD(P)-linked oxidoreductase superfamily protein

Dcitr01g22340.1.1	-8.455	0.000	Unknown protein
