## Supplementary material for "Quantification of new and archived *Diaphorina citri* transcriptome data using a chromosomal length *D. citri* genome assembly reveals the vector’s tissue-specific transcriptional response to citrus greening disease": Table S5

**Table S5:** All transcripts from the bacteriome dataset that have differential expression log2FoldChange>|2| and adjusted p-value<0.05. Sorted by Log2FoldChange.

**Transcript_ID	L2FC	padj	Annotation**

Dcitr01g16790.1.1	8.432	0.000	Cationic amino acid transporter

Dcitr04g03620.1.1	7.487	0.000	Methyltransferase family protein

Dcitr01g21740.1.1	7.335	0.000	Thioredoxin domain-containing protein 17

Dcitr06g12720.1.2	6.793	0.001	Sorbin and SH3 domain-containing protein 2

Dcitr01g12970.1.1	6.466	0.004	Zinc-binding dehydrogenase

Dcitr10g11050.1.1	6.295	0.001	Unknown protein

Dcitr01g09990.1.1	6.114	0.006	Androgen-induced gene 1 protein

Dcitr08g01750.1.2	5.977	0.007	Blood vessel epicardial substance

Dcitr07g08590.1.1	5.592	0.028	Major facilitator transporter

Dcitr04g12240.1.1	5.483	0.006	Lactoylglutathione lyase

Dcitr06g04730.1.1	5.482	0.025	Phthiotriol/phenolphthiotriol dimycocerosates methyltransferase

Dcitr02g04200.1.2	5.314	0.005	Dystroglycan

Dcitr11g06040.1.1	5.140	0.011	GATA zinc finger domain-containing protein 10-like, partial

Dcitr13g03670.1.1	5.090	0.012	Gag-Pro-Pol polyprotein

Dcitr00g02740.1.1	4.974	0.020	RING/U-box

Dcitr03g07550.1.1	4.541	0.004	Delta-1-pyrroline-5-carboxylate synthase

Dcitr04g04380.1.1	4.539	0.022	Piezo-type mechanosensitive ion channel component 1

Dcitr01g05990.1.1	4.446	0.002	Acid phosphatase-like protein 2

Dcitr00g02000.1.1	4.360	0.025	Unknown

Dcitr10g03040.1.3	4.353	0.000	SH3 domain-containing protein Dlish

Dcitr04g01760.1.1	4.304	0.026	LOW QUALITY PROTEIN: uncharacterized protein LOC112591404

Dcitr00g11870.1.1	4.210	0.007	Elongation

Dcitr01g20610.1.1	4.122	0.011	F-actin-capping protein subunit alpha

Dcitr04g01700.1.1	4.066	0.005	Erg28-domain containing protein

Dcitr06g09230.1.1	4.053	0.005	COP9 signalosome complex subunit 7a

Dcitr03g07170.1.1	4.034	0.008	60S ribosomal protein L26

Dcitr05g16270.1.1	3.932	0.027	early endosome antigen 1-like

Dcitr12g10310.1.1	3.875	0.050	Unknown protein

Dcitr08g11110.1.1	3.744	0.007	Agrin

Dcitr03g19430.1.1	3.733	0.021	Hexokinase

Dcitr01g18830.1.1	3.609	0.024	sphingomyelin phosphodiesterase 4-like

Dcitr02g02510.1.1	3.555	0.008	Phosphoenolpyruvate synthase

Dcitr02g19280.1.1	3.319	0.019	Poly [ADP-ribose] polymerase

Dcitr00g12300.1.1	3.299	0.020	Protein

Dcitr01g18190.1.1	3.254	0.032	60S ribosomal protein L37a

Dcitr04g05130.1.1	3.212	0.039	Unknown protein

Dcitr13g06110.1.1	3.158	0.040	Unknown protein

Dcitr05g10980.1.1	3.122	0.016	RING finger and SPRY domain-containing protein 1

Dcitr03g08980.1.1	3.075	0.000	Unknown protein

Dcitr07g01070.1.1	3.071	0.002	L-allo-threonine aldolase

Dcitr11g01560.1.1	3.070	0.030	Cation-chloride cotransporter 1

Dcitr02g19260.1.1	2.937	0.014	Ribosomal protein L23

Dcitr07g01080.1.2	2.899	0.025	Myosin heavy chain, non-muscle

Dcitr03g19060.1.1	2.863	0.000	protein phosphatase 1 regulatory subunit 21-like

Dcitr12g05400.1.1	2.853	0.024	5'-nucleotidase

Dcitr03g08990.1.1	2.549	0.000	Unknown protein

Dcitr01g01470.1.1	2.540	0.006	Poly(A) RNA polymerase, mitochondrial

Dcitr06g08650.1.1	2.535	0.000	Lipase

Dcitr05g01800.1.1	2.473	0.000	PiggyBac transposable element-derived protein 4

Dcitr11g09370.1.1	2.456	0.038	Kanadaptin

Dcitr04g11810.1.1	2.446	0.000	Transposon Ty3-I Gag-Pol polyprotein

Dcitr04g09800.1.1	2.413	0.036	Aldose 1-epimerase

Dcitr04g01290.1.1	2.392	0.000	Mortality factor 4-like protein 1

Dcitr05g14050.1.1	2.344	0.011	Cytochrome P450 305E1

Dcitr13g05490.1.1	2.269	0.000	Aldehyde dehydrogenase

Dcitr04g01240.1.1	2.268	0.028	Mitotic checkpoint serine/threonine-protein kinase BUB1

Dcitr05g07320.1.1	2.260	0.030	N-acetylaspartate synthetase

Dcitr05g06560.1.1	2.216	0.004	LETM1 domain-containing protein 1

Dcitr02g04730.1.1	2.207	0.000	Spondin-1

Dcitr07g01110.1.1	2.172	0.000	Myosin heavy chain, non-muscle

Dcitr04g11820.1.1	2.162	0.000	Transposon Ty3-I Gag-Pol polyprotein

Dcitr05g17180.1.1	2.136	0.013	DnaJ subfamily C member 22

Dcitr03g19600.1.1	2.106	0.007	Protein arginine N-methyltransferase

Dcitr03g09890.1.1	2.072	0.035	Unknown protein

Dcitr04g11220.1.2	2.069	0.003	pre-mRNA-splicing factor CWC22 homolog isoform X2

Dcitr05g15210.1.1	2.058	0.035	serine/threonine-protein phosphatase 6 regulatory subunit 3-A

Dcitr11g01990.1.1	2.052	0.001	Pro-Pol polyprotein

Dcitr09g07400.1.1	2.039	0.036	Ribonuclease

Dcitr00g12410.1.1	2.035	0.047	E3

Dcitr13g01090.1.1	2.013	0.000	Myb

Dcitr01g09300.1.1	-2.059	0.004	Unknown protein

Dcitr02g18210.1.1	-2.136	0.037	ABC transporter G family

Dcitr03g02070.1.1	-2.147	0.022	Fibroblast growth factor 17

Dcitr03g18480.1.1	-2.176	0.038	Histone acetyltransferase type B catalytic subunit

Dcitr11g03410.1.1	-2.302	0.018	Unconventional myosin-IXa

Dcitr12g03860.1.1	-2.316	0.022	organic solute transporter ostalpha protein (DUF300) Chr5:9292436-9294407 FORWARD LENGTH%3D422 201606

Dcitr07g08190.1.1	-2.318	0.009	Peroxisomal N(1)-acetyl-spermine/spermidine oxidase

Dcitr11g06580.1.1	-2.341	0.024	CLUMA_CG018496, isoform A

Dcitr07g06540.1.1	-2.361	0.000	4-hydroxy-tetrahydrodipicolinate synthase

Dcitr10g02220.1.1	-2.381	0.014	Cytochrome c oxidase subunit

Dcitr04g06980.1.1	-2.412	0.000	E3 ubiquitin-protein ligase E3D

Dcitr00g06360.1.1	-2.513	0.040	Sorting

Dcitr08g10240.1.1	-2.520	0.022	Filamin-B

Dcitr00g10370.1.1	-2.576	0.006	Sphingomyelin

Dcitr03g07570.1.1	-2.605	0.037	Delta-1-pyrroline-5-carboxylate synthase

Dcitr02g04720.1.1	-2.660	0.000	DNA helicase

Dcitr05g15920.1.1	-2.794	0.004	Major facilitator superfamily domain-containing protein 6

Dcitr02g10750.1.1	-2.841	0.016	minichromosome maintenance domain-containing protein 2 isoform X1

Dcitr07g03820.1.1	-2.848	0.006	Aminopeptidase

Dcitr01g17200.1.1	-2.885	0.012	Prolyl oligopeptidase family protein

Dcitr03g21220.1.1	-2.934	0.002	hemicentin-1-like

Dcitr01g05380.1.1	-3.088	0.035	transcription factor A, mitochondrial

Dcitr07g09000.1.1	-3.272	0.038	Alkaline nuclease

Dcitr00g06180.1.1	-3.618	0.041	Broad-complex

Dcitr01g06690.1.1	-3.653	0.003	Elongation factor 4

Dcitr04g07450.1.1	-3.702	0.008	retinaldehyde-binding protein 1-like

Dcitr03g09640.1.1	-3.744	0.007	Unknown protein

Dcitr06g06000.1.1	-3.760	0.000	Ankyrin repeat family protein

Dcitr03g01060.1.1	-3.764	0.005	L-lactate dehydrogenase

Dcitr00g09080.1.1	-3.788	0.007	Krueppel-like

Dcitr09g04720.1.2	-3.967	0.008	Vacuolar

Dcitr12g08510.1.4	-3.982	0.000	Bax inhibitor 1-related

Dcitr08g10290.1.1	-4.023	0.047	HEAT repeat-containing protein 5B

Dcitr04g11300.1.1	-4.444	0.044	Pre-mRNA-splicing factor CWC22

Dcitr07g06450.1.1	-4.566	0.024	2-oxoglutarate (2OG) and Fe(II)-dependent oxygenase superfamily protein

Dcitr03g03100.1.1	-4.640	0.019	regulator of G-protein signaling loco-like

Dcitr04g07640.1.1	-4.648	0.003	Protein SYS1 homolog

Dcitr06g14870.1.1	-4.749	0.027	Coiled-coil domain-containing protein 149

Dcitr01g22730.1.1	-4.786	0.000	RING/U-box superfamily protein Chr3:5102210-5104082 REVERSE LENGTH%3D249 201606

Dcitr10g04230.1.1	-4.940	0.020	CAP-Gly domain-containing linker protein 1

Dcitr03g19580.1.1	-5.409	0.050	Delta-aminolevulinic acid dehydratase

Dcitr06g03080.1.1	-5.828	0.000	Unknown protein

Dcitr10g04540.1.1	-6.068	0.024	NADPH--cytochrome P450 reductase
