## Supplementary material for "Quantification of new and archived *Diaphorina citri* transcriptome data using a chromosomal length *D. citri* genome assembly reveals the vector’s tissue-specific transcriptional response to citrus greening disease": Table S6

**Table S6:** All transcripts from the head dataset that have differential expression log2FoldChange>|2| and adjusted p-value<0.05. Sorted by Log2FoldChange.

| **HEAD** | **padj** | **L2FC** | **Annotation** |
| --- | --- | --- | --- |
| Dcitr05g09170.1.1 | 0.039 | -5.333 | ATP synthase delta subunit |
| Dcitr05g06500.1.1 | 0.032 | -4.263 | Unknown protein |
| Dcitr02g01080.1.1 | 0.027 | -4.095 | Vigilin |
| Dcitr01g21980.1.1 | 0.027 | -3.426 | RNA-directed DNA polymerase from mobile element jockey |
| Dcitr02g09580.1.1 | 0.032 | 3.415 | intracellular protein transport protein USO1 |
| Dcitr04g10550.1.1 | 0.001 | 5.276 | neuromodulin |
| Dcitr07g08870.1.1 | 0.024 | 5.289 | Rho GTPase-activating protein 17 |
| Dcitr10g06500.1.1 | 0.005 | 5.562 | Glucocorticoid-induced transcript 1 protein |
| Dcitr05g04030.1.1 | 0.043 | 5.615 | Tumor protein p53-inducible nuclear protein 1 |
| Dcitr08g09760.1.3 | 0.014 | 5.633 | Insulin-like growth factor 2 mRNA-binding protein 3 |
