## Supplementary material for "Quantification of new and archived *Diaphorina citri* transcriptome data using a chromosomal length *D. citri* genome assembly reveals the vector’s tissue-specific transcriptional response to citrus greening disease": Table S7

**Table S7:** Data used to generate Figure 3, including annotations, p-values and Log2FoldChange values for each transcript listed.

| Endocytosis |  |  |  |  |  |  |
| --- | --- | --- | --- | --- | --- | --- |
| Gene_ID | Midgut | padj | Bacteriome | padj | Salivary Gland | padj |
| Dcitr01g20610.1.1 | 0.04 | 0.99 | 4.12 | 0.01 | -2.79 | 0.14 |
| Dcitr02g07360.1.4 | -7.23 | 0.00 | -0.21 | 0.99 | NA | NA |
| Dcitr05g12140.1.1 | 3.75 | 0.04 | -0.51 | 0.79 | 0.56 | 0.97 |
| Dcitr05g16270.1.1 | 0.88 | 0.58 | 3.93 | 0.03 | NA | NA |
| Dcitr07g01080.1.2 | -0.77 | NA | 2.90 | 0.03 | 4.62 | 0.15 |
| Dcitr07g01110.1.1 | -2.44 | 0.49 | 2.17 | 0.00 | 2.01 | 0.65 |
| Dcitr07g08110.1.1 | 4.32 | 0.03 | -0.69 | NA | NA | NA |
| Dcitr09g04720.1.2 | -0.44 | NA | -3.97 | 0.01 | 0.27 | 0.98 |
| Dcitr11g04950.1.2 | 3.97 | 0.00 | 0.12 | 1.00 | 3.25 | 0.20 |
| Dcitr12g03760.1.1 | 3.62 | 0.02 | -1.05 | 0.96 | 1.29 | 0.95 |
| Dcitr12g03760.1.1 | 3.62 | 0.01 | -1.05 | 0.58 | 1.29 | 0.00 |
| Dcitr12g07070.1.1 | NA | 0.00 | 0.88 | 0.87 | 2.53 | 0.99 |
| Dcitr13g01820.1.1 | 3.60 | 0.02 | -0.57 | 0.96 | -1.17 | 0.95 |
| Dcitr13g01840.1.1 | 2.51 | NA | 2.82 | 0.19 | 5.78 | 0.01 |
| Dcitr13g01890.1.1 | 2.96 | 0.02 | -1.05 | 0.04 | 0.52 | 0.88 |
| Ubiquination |  |  |  |  |  |  |
| Gene_ID | Midgut | padj | Bacteriome | padj | Salivary Gland | padj |
| Dcitr00g12410.1.1 | 0.60 | 0.84 | 2.04 | 0.05 | 0.64 | 0.98 |
| Dcitr01g04810.1.1 | -4.39 | 0.00 | -1.11 | 0.00 | -0.71 | 0.98 |
| Dcitr01g10860.1.1 | -2.83 | 0.04 | 0.13 | 0.98 | 0.65 | 0.98 |
| Dcitr01g22250.1.1 | 4.76 | 0.01 | 2.31 | NA | 0.71 | 0.98 |
| Dcitr02g02390.1.2 | 0.05 | 1.00 | 0.18 | 1.00 | -5.32 | 0.01 |
| Dcitr02g09710.1.1 | 4.57 | 0.00 | -0.53 | 0.99 | 0.42 | 0.98 |
| Dcitr04g06980.1.1 | -0.01 | 1.00 | -2.41 | 0.00 | 0.57 | 0.98 |
| Dcitr04g09440.1.1 | -1.94 | 0.54 | 0.08 | 0.99 | 11.36 | 0.00 |
| Dcitr06g04060.1.1 | -1.52 | 0.52 | 2.86 | 0.57 | 4.69 | 0.03 |
| Dcitr10g09310.1.1 | 3.43 | 0.05 | -0.91 | 0.86 | 2.08 | 0.49 |
| Dcitr13g03480.1.1 | 0.24 | 0.93 | -0.23 | 0.78 | 7.41 | 0.00 |
| Dcitr13g04800.1.1 | 0.29 | 0.95 | -1.93 | NA | 6.05 | 0.00 |
| Immunity |  |  |  |  |  |  |
| Gene_ID | Midgut | padj | Bacteriome | padj | Salivary Gland | padj |
| Dcitr01g11640.1.1 | -0.07 | 0.99 | -3.66 | NA | -6.26 | 0.02 |
| Dcitr02g03230.1.1 | 4.03 | 0.00 | 0.18 | 0.99 | 1.28 | 0.62 |
| Dcitr02g04530.1.1 | 0.32 | 0.91 | -0.65 | 0.16 | 3.70 | 0.01 |
| Dcitr02g04730.1.1 | 2.53 | 0.38 | 2.21 | 0.00 | 10.13 | 0.00 |
| Dcitr03g07590.1.1 | 3.51 | 0.00 | -0.80 | 0.95 | 5.79 | 0.02 |
| Dcitr03g15730.1.1 | 2.66 | 0.15 | -1.75 | 0.85 | 0.15 | 0.99 |
| Dcitr03g15750.1.1 | 2.18 | 0.41 | -1.95 | 0.81 | 4.46 | 0.01 |
| Dcitr03g15760.1.1 | 3.02 | 0.01 | -0.90 | 0.94 | 2.29 | 0.36 |
| Dcitr03g15900.1.1 | -2.26 | 0.01 | -0.47 | 0.72 | 1.21 | 0.95 |
| Dcitr07g03820.1.1 | -1.58 | 0.73 | -2.85 | 0.01 | 3.65 | 0.01 |
| Dcitr07g08110.1.1 | 4.32 | 0.03 | -0.69 | NA | 5.86 | 0.01 |
| Dcitr10g09840.1.1 | 1.44 | 0.41 | -0.61 | 0.98 | -0.14 | 0.99 |
| Dcitr05g07530.1.1 | NA | NA | -0.14 | 0.99 | 0.06 | 1.00 |
| Dcitr09g07400.1.1 | -0.31 | 0.98 | 2.04 | 0.04 | 0.67 | 0.98 |
| Dcitr10g06970.1.2 | -0.71 | 0.94 | 0.14 | 0.99 | NA | NA |
| Dcitr10g10180.1.1 | 0.80 | 0.65 | -0.66 | 0.45 | 5.17 | 0.04 |
| Ribosomal |  |  |  |  |  |  |
| Gene_ID | Midgut | padj | Bacteriome | padj | Salivary Gland | padj |
| Dcitr01g11490.1.1 | -4.51 | 0.00 | -0.13 | 0.96 | NA | NA |
| Dcitr01g18190.1.1 | -2.82 | 0.37 | 3.25 | 0.03 | 0.22 | 0.99 |
| Dcitr02g04500.1.1 | -0.52 | 0.94 | NA | NA | 4.69 | 0.03 |
| Dcitr02g14920.1.1 | 3.40 | 0.00 | -1.09 | 0.40 | -0.68 | 0.98 |
| Dcitr02g19260.1.1 | -0.25 | NA | 2.94 | 0.01 | 2.25 | 0.52 |
| Dcitr02g19280.1.1 | -0.42 | 0.62 | 3.32 | 0.02 | 0.33 | 0.99 |
| Dcitr03g04260.1.1 | -2.60 | NA | 0.30 | 0.99 | 5.77 | 0.00 |
| Dcitr03g04400.1.1 | 2.75 | NA | -0.27 | 0.99 | 5.53 | 0.00 |
| Dcitr03g07170.1.1 | 1.79 | 0.60 | 4.03 | 0.01 | 0.65 | 0.98 |
| Dcitr04g05810.1.1 | 6.17 | 0.00 | 0.25 | 0.98 | -2.62 | 0.77 |
| Dcitr04g12470.1.1 | -4.48 | 0.00 | -0.70 | 0.89 | -0.83 | 0.88 |
| Dcitr04g13690.1.1 | NA | NA | -0.09 | 1.00 | 10.52 | 0.00 |
