## Supplementary material for "Quantification of new and archived *Diaphorina citri* transcriptome data using a chromosomal length *D. citri* genome assembly reveals the vector’s tissue-specific transcriptional response to citrus greening disease": Table S8

**Table S8:** All transcripts from the salivary gland dataset that have predicted signal sequences. The four in **bold** text had predicted transmembrane helices.

| **Transcript ID** | **padj** | **L2FC** | **Annotation** | **Mw (kD)** | **Cleavage site** | **AA site** |
| --- | --- | --- | --- | --- | --- | --- |
| Dcitr05g04310.1.1 | 0.003 | 5.399 | AGAP013432-PA | 10.12 | 14-15 | ALC-DQ |
| Dcitr03g16830.1.3 | 0.000 | 7.171 | basic salivary proline-rich protein 3-like | 14.17 | 25-26 | VIG-QS |
| Dcitr03g15730.1.1 | 0.010 | 3.646 | CLIPB - Serine Protease 1 | 35.39 | 22-23 | GLA-YS |
| **Dcitr09g07790.1.1** | **0.045** | **-4.641** | **conserved** | **57.12** | **20-21** | **ISA-ES** |
| **Dcitr13g04790.1.1** | **0.020** | **2.952** | **Lipid phosphate phosphatase** | **34.06** | **21-22** | **SKQ-GF** |
| Dcitr07g06610.1.1 | 0.000 | 5.910 | Meteorin-like protein | 36.49 | 28-29 | ITG-LV |
| Dcitr03g15750.1.1 | 0.011 | 5.861 | Serine protease | 35.19 | 22-23 | GLA-YS |
| **Dcitr07g08050.1.1** | **0.046** | **3.944** | **Transmembrane emp24 domain-containing protein 7** | **25.58** | **28-29** | **VQA-VE** |
| **Dcitr03g13400.1.2** | **0.000** | **-4.409** | **UDP-glucuronosyltransferase** | **57.55** | **19-20** | **AQG-AN** |
| Dcitr01g04160.1.1 | 0.000 | 5.462 | Unknown protein | 29.65 | 19-20 | TQS-QL |
| Dcitr05g09660.1.1 | 0.030 | 4.854 | Unknown protein | 6.87 | 21-22 | TLA-SD |
| Dcitr01g22340.1.1 | 0.000 | -8.455 | Unknown protein | 75.36 | 20-21 | TLC-GV |
