## Supplementary material for "Quantification of new and archived *Diaphorina citri* transcriptome data using a chromosomal length *D. citri* genome assembly reveals the vector’s tissue-specific transcriptional response to citrus greening disease": Table S9

**Table S9:** All piggyBac-related genes currently annotated in the Diaci_v3.0 genome. The gene identified in our transcript analysis is in **bold**.

| **Gene_ID** | **Annotation** |
| --- | --- |
| Dcitr02g01360.1.1 | PiggyBac transposable element-derived protein 4 |
| Dcitr03g11500.1.1 | PiggyBac transposable element-derived protein 2 |
| **Dcitr05g01800.1.1** | **PiggyBac transposable element-derived protein 4** |
| Dcitr05g01980.1.1 | PiggyBac transposable element-derived protein 4 |
| Dcitr05g02000.1.1 | PiggyBac transposable element-derived protein 4 |
| Dcitr08g12690.1.1 | PiggyBac transposable element-derived protein 2 |
| Dcitr09g02860.1.1 | PiggyBac |
| Dcitr09g03340.1.1 | PiggyBac |
| Dcitr10g03650.1.1 | piggyBac transposable element-derived protein 3-like |
| Dcitr10g10330.1.1 | PiggyBac transposable element-derived protein 2 |
| Dcitr11g09800.1.1 | PiggyBac transposable element-derived protein 4 |
